# APOE^ε4^ and exercise interact to influence systemic and cerebral risk factors for dementia

**DOI:** 10.1101/2022.03.01.480612

**Authors:** Kate E. Foley, Cory A. Diemler, Amanda A. Hewes, Dylan T. Garceau, Michael Sasner, Gareth R. Howell

## Abstract

**INTRODUCTION:** *APOE^ε4^* is the strongest genetic risk factor for Alzheimer’s disease and related dementias (ADRDs) affecting many different pathways that lead to cognitive decline. Exercise is one of the most widely proposed prevention, and intervention strategies to mitigate risk and symptomology of ADRDs. Importantly, exercise and *APOE^ε4^* affect similar processes on the body and brain. While both *APOE^ε4^*, and exercise have been studied extensively, their interactive effects are not well understood.

**METHODS:** To address this, male and female *APOE^ε3/ε3^*, *APOE^ε3/ε4^* and *APOE^ε4/ε4^* mice ran voluntarily from wean (1mo) to midlife (12mo). Longitudinal and cross-sectional phenotyping was performed on the periphery and the brain, on markers of risk for dementia such as weight, body composition, circulating cholesterol composition, activities of daily living, energy expenditure, and cortical and hippocampal transcriptional profiling.

**RESULTS:** Data revealed chronic running decreased age-dependent weight gain, lean and fat mass, and serum LDL concentration dependent on *APOE* genotype. Additionally, murine activities of daily living and energy expenditure were significantly influenced by an interaction between *APOE* genotype and running in both sexes. Transcriptional profiling of the cortex and hippocampus predicted that *APOE* genotype and running interact to affect numerous biological processes including vascular integrity, synaptic/neuronal health, cell motility, and mitochondrial metabolism, in a sex-specific manner.

**DISCUSSION:** These data provide compelling evidence that *APOE* genotype should be considered for population-based strategies that incorporate exercise to prevent ADRDs.

## 1. Background

Aging and *APOE^ε4^* are the strongest risk factors for Alzheimer’s disease and related dementias (ADRDs)[1]. With *APOE^ε4^* implicated in unfavorable systemic changes such as high BMI, dysregulated cholesterol concentrations, and aberrant metabolism, as well as deficits in cerebral health such as changes in cerebral metabolism, cerebrovasculature, and neuronal health, the *APOE^ε4^* allele has been targeted to help reverse these risks[2–7]. The cerebral changes caused by *APOE^ε4^* emerge in humans at early ages and can worsen with advancing age[8–12]. Further, the impact of *APOE^ε4^* dosage (such as in the *APOE^ε3/ε4^* versus *APOE^ε4/ε4^* genotype) on peripheral and brain health during aging is understudied. Targeting *APOE^ε4^* through pharmacological interventions has resulted in both beneficial and damaging outcomes meaning therapies targeting APOE-dependent pathways will likely need to be tailored to specific mechanisms[13–16].

While pharmacological interventions are still being investigated, others have turned to non-pharmacological interventions to reduce risk for ADRDs, such as exercise[13, 17]. Studies in mice show benefits of exercise to peripheral health, as well as improvements to cognitive function[18–27]. Though the cognitive changes due to exercise have been controversial, with human studies showing either no change or improvements with exercise, it is widely accepted that exercise affects the body in a generally positive manner (i.e., decreasing weight/fat mass, improving metabolism and circulation, and elevating mood)[19, 28–33]. While understanding the effect of exercise on neuronal health is critical, other compartments of the brain are largely neglected. It is essential to understand how exercise affects all mechanisms that pertain to ADRD risk, such as metabolism and vascular health.

It is unknown if the detrimental effects associated with *APOE^ε4^* can be mitigated by exercise, or conversely, whether the effects of exercise are impacted by *APOE^ε4^* genotype. Studies in humans are performed later in life after symptom onset, typically measuring improvements to activities of daily living and quality of life. While important, it is necessary to understand whether running can influence risk factors for dementia before symptomology. We evaluated the systemic and cerebral effects of running across *APOE^ε3/ε3^*, *APOE^ε3/ε4^* and *APOE^ε4/ε4^* litter-matched mice during early aging. We show that chronic running affects multiple ADRD-relevant phenotypes in both the periphery and the brain but these effects are both *APOE* genotype- and sex-specific.

## 2. Methods

### 2.1 Mouse Husbandry

Novel *APOE* mouse strains were created on C57BL/6J (B6) and maintained at The Jackson Laboratory as previously described prior[34]. Mice were kept in a 12/12-hour light/dark cycle (06:00 – 18:00 light) and fed ad libitum 6% kcal fat standard mouse chow. Experimental cohorts were generated by intercrossing male and female *APOE^ε3/ε4^* mice to create *APOE^ε3/ε3^*, *APOE^ε3/ε4^* and *APOE^ε4/ε4^* male and female littermate controls. Animals were divided as evenly as possible per litter into running and sedentary cohorts. All experiments were approved by the Institutional Animal Care and Use Committee (IACUC) at The Jackson Laboratory.

### 2.2 Exercise by Voluntary Running

Mice were group-housed into two or three per pen and given 24-hour access to an unlocked (running) or locked (sedentary) running wheel (Innovive Inc). At 5 months (5mo), mice were singly housed for the duration of the experiment. At 6 and 11mo, running mice were tracked for number of rotations per minute for five to seven nights during the dark cycle when they are most active using trackable running wheels (Med Associates Inc.). Any nights that had fewer than 700 minutes tracked were excluded from analysis. For each mouse, sum of rotations per night was calculated and then averaged across all nights.

### 2.3 Harvesting, Tissue Preparation, Plasma Collection

All mice were euthanized by intraperitoneal injection of a lethal dose of Ketamine (100mg/ml)/Xylazine(20mg/ml). Mice were perfused intracardially with 1XPBS. Brains were carefully dissected, hemisected sagittally, and one half was then snap frozen on solid CO_2_ for later dissection and RNA-sequencing. At timepoints throughout the experiment, blood plasma was collected via cheek bleed. Blood was carefully collected in K2 EDTA (1.0mg) microtainer tubes (BD), allowed to sit at room temperature for at least 30 minutes, and then centrifuged at 21°C for 10 minutes at 5000rpm. Plasma was carefully collected and stored at −20°C. At the harvest timepoint (12mo), blood was collected in K2 EDTA (1.0mg) microtainer tubes (BD) through cardiac puncture. Plasma total cholesterol (mg/dL), direct LDL (mg/dL), and HDL (mg/dL) concentrations were characterized on the Beckman Coulter AU680 chemistry analyzer. All samples were profiled at the same time at the end of the experiment to avoid batch effects.

### 2.4 Nuclear Magnetic Resonance Imaging (NMR)

Each cohort was subjected to NMR imaging at 6 and 11mo. NMR was performed as previously described[35]. Briefly, weight was measured, and mice were briefly placed into a Plexiglas tube 2.5 in. by 8 in. which was then subjected to NMR (EchoMRI, Houston, TX). Magnetic field was measured by a 5-gauss magnet. Measurements included weight, lean muscle mass, and fat mass, as well as fat percentage ((fat/body weight) × 100).

### 2.6 Activities of daily living and Indirect Calorimetry

After NMR measurements, groups of 16 mice were measured at a time for energy balance through indirect calorimetry measurement cages (Sable Promethion). Briefly, these specialized cages continuously measure food and water intake, general activity (pedometers), wheel running behavior, energy expenditure (kcal/hr), and respiratory quotient (RQ). Measurements are collected for five days in five-minute interval bins. The respiratory quotient (RQ) is a ratio of the volume of carbon dioxide (CO_2_) released over the volume of oxygen (O_2_) absorbed. RQ has been widely used in humans and mice as a tool to determine the starting substrate for energy metabolism (carbohydrate RQ ~1, protein RQ ~0.8, fat RQ ~0.7, anaerobic respiration RQ ~0, and multiple energy sources RQ ~0.8) [36–41].

### 2.7 RNA-sequencing, linear modeling and GSEA

RNA extraction, library construction, RNA sequencing and seq quality control was performed as described previously[34, 35]. Genes were then filtered by 1) removing all genes that did not vary in expression (gene count change across all samples was 0) and 2) removing all genes that did not have at least five reads in 50% of the samples. Remaining genes (20,641) were normalized using DEseq2[42]. Principal component analysis (PCA) on the variance stabilized data (vst) identified outliers. To allow for the evaluation of *APOE^ε4^* allele dosage, each linear model included two genotype comparisons: 1) *APOE^ε3/ε4^* to *APOE^ε3/ε3^* and 2) *APOE^ε4/ε4^* to *APOE^ε3/ε3^*. Linear models were run separately for 1) cortex - female, 2) cortex – male, 3) hippocampus – female, and 4) hippocampus – male. β-estimates were obtained for all four linear models that evaluated the main effects of *APOE* genotype (*APOE^ε3/ε4^*, *APOE^ε4/ε4^*, ref: *APOE^ε3/ε3^*) and running (run, ref: sed), as well as the interaction between *APOE* genotype and running (*APOE^ε3/ε4^*:Run, *APOE^ε4/ε4^*:Run). For each linear model, gseGO from the clusterProfiler package was run on genes significant for each factor. Gene Set Enrichment Analysis (GSEA) was used to determine GO terms for the genes significant for the main (running, *APOE^ε3/ε4^*, *APOE^ε4/ε4^*) and interacting factors (*APOE^ε3/ε4^*:Run and *APOE^ε4/ε4^*:Run). Normalized Enrichment Scores (NES) from GSEA were used to identify terms that were positively or negatively associated with each factor. GO terms were ordered based on the NES. Terms with a positive NES had more genes higher on the ranked list (ie. more positive β values) and the terms with a negative NES containing more genes lower on the ranked list (ie., more negative β values). Enriched GO terms had overlapping biological functions that we termed ‘vascular integrity’, ‘cellular motility’, ‘immune system response’, ‘mitochondrial metabolism’, and ‘synaptic/neuronal health’. The top 20 most positive and negative GO terms were visualized for the cortex and hippocampus for both females and males (**Supp Figs 10-24**).

### 2.10 Statistical Analysis

For all weights and body composition analysis, a two-way ANOVA for *APOE* genotype, activity, and the interaction between *APOE* genotype and activity was calculated. Bonferroni post hoc corrections were calculated and significance within genotype (the effect of running per genotype) was visualized.

## 3. Results

### *APOE* genotype did not affect voluntary running from young to midlife

To determine the effects of one *APOE^ε4^* allele to two *APOE^ε4^* alleles, we compared the *APOE^ε3/ε4^* and *APOE^ε4/ε4^* genotypes to the control, *APOE^ε3/ε3^* genotype (**Fig 1A**). Previous studies have shown that females run more than males, therefore we assessed the sexes separately[35]. There was no difference in voluntary running during the dark cycle across *APOE* genotypes, however there was expected variation between individual mice within the *APOE* genotypes (**Fig 1B-C**, **Supp Fig 1-2**). There was an age-dependent decrease in voluntary running from 6-11mo, however there was no difference between *APOE* genotypes (**Fig 1D-G**). These findings show that running is not a variable between the *APOE* genotypes, and therefore not a confound in subsequent analyses.

**Figure 1:**
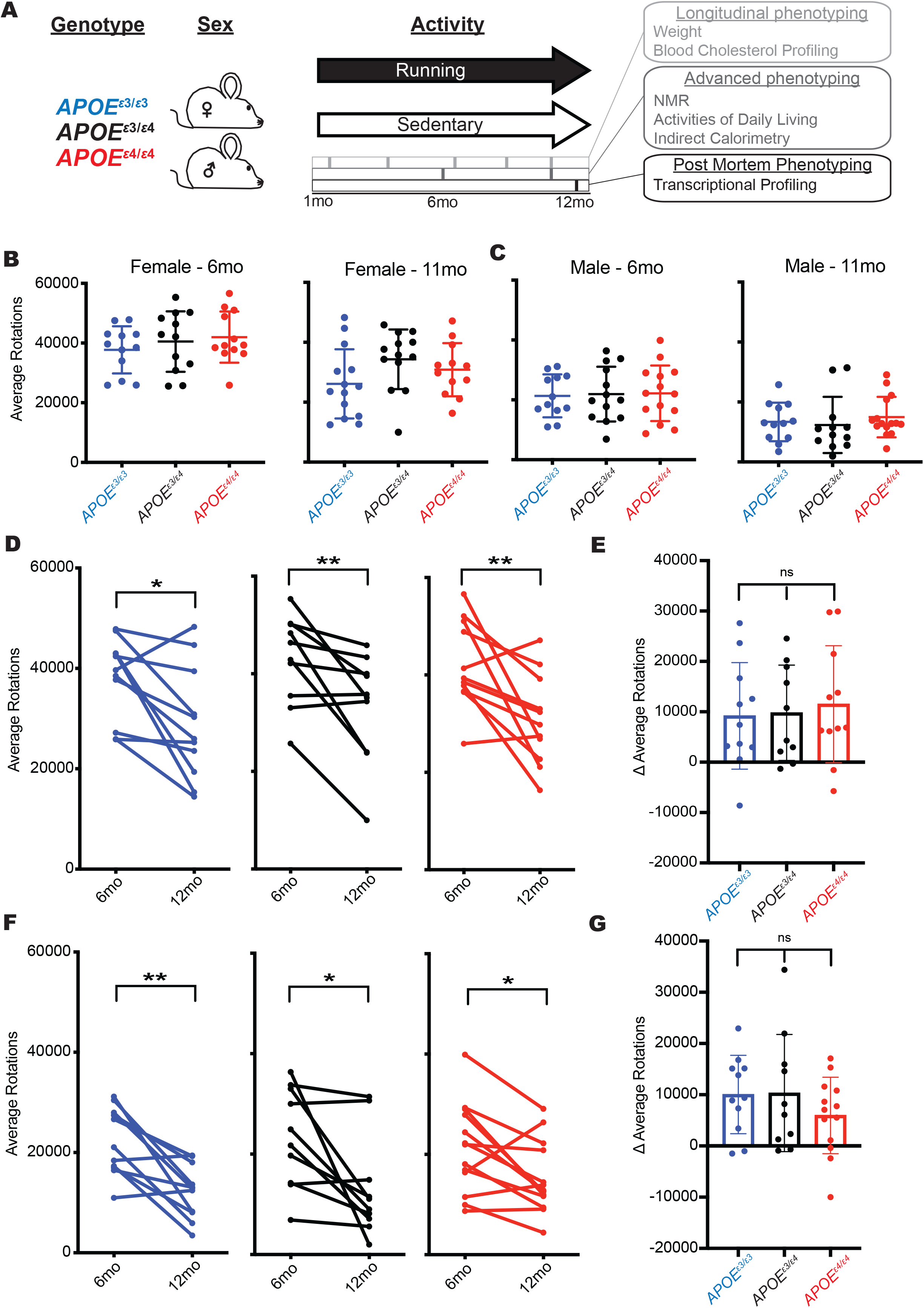
Voluntary chronic running to midlife is not different between *APOE* genotypes. **(A)** Schematic of the voluntary running paradigm where *APOE^ε3/ε3^*, *APOE^ε3/ε4^* and *APOE^ε4/ε4^* male and female mice were introduced to a locked (control – sedentary) or unlocked (treatment - running) running wheel at 1 month (mo) until 12mo (midlife). Longitudinal, advanced, and post-mortem phenotyping is indicated. (**B-C)** No difference in running (average rotations across multiple consecutive nights) between *APOE* genotypes at both six and eleven months in female (B) or male (C) mice. **(D-G)** Average rotations per mouse at 6mo and 12mo showed an age-dependent decrease for both females (D) and males (F) however the change over time was not significantly different between genotypes **(E,G)**. Data presented as mean ± SD, one way ANOVA with Tukey’s multiple comparison performed for **B,C,E,G**. Two-sided paired t-test performed for **D,F**. *P < 0.05, **P < 0.01.

### *APOE* genotype and running interact in a sex-specific manner to modulate general markers of healthy aging

Weight, body composition (e.g., lean mass, fat mass, and fat percentage) and cholesterol levels are commonly used as a general proxy for health in humans[43–45]. These biometrics are typically measured at routine physicals and are considered indicative of general health status, and markers for obesity, cardiovascular disruption, and lipid dysregulation[46–49]. We examined whether running affected weight, body composition and cholesterol across *APOE* genotypes. Monthly weights (from 1-12mos) revealed an expected age-dependent weight gain in sedentary mice that was significantly attenuated by running (**Fig 2AF, Supp Fig 3A-D**). In females, but not males, the *APOE^ε4/ε4^* genotype caused a greater running-based attenuation in weight gain compared to *APOE^ε3/ε3^* and *APOE^ε3/ε4^*. These results suggest that the beneficial effects of running on weight loss are *APOE* genotype-dependent in females only.

**Figure 2:**
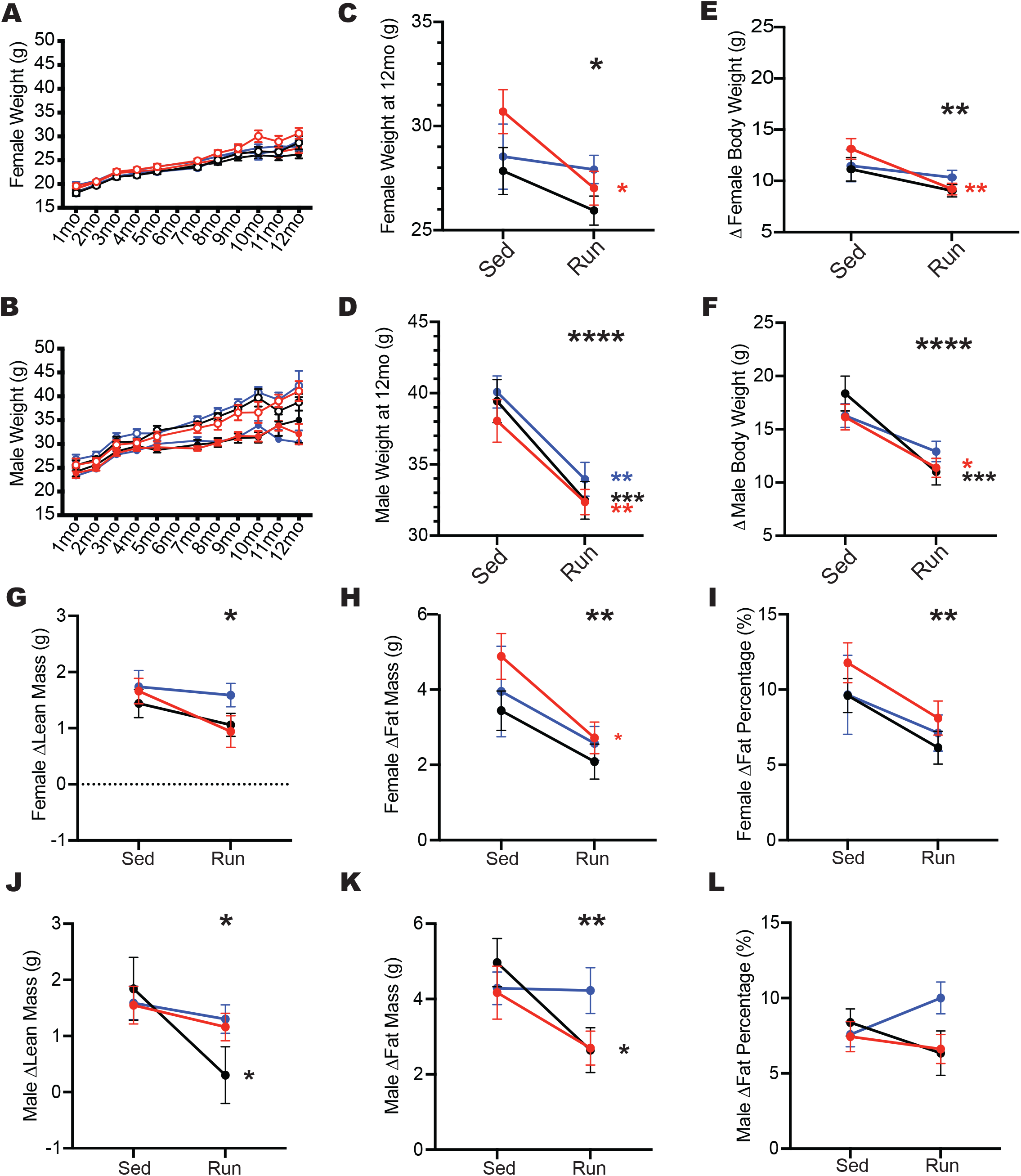
Running attenuated age-dependent weight gain and fat accumulation across *APOE* genotypes. **(A-B)** Expected age-dependent weight gain from one to twelve months in females (A) and males (B). **(C-D)** Running mice weighed significantly less at 12mo in both females (C) and males (D). **(E-F)** Running significantly attenuated age-dependent weight gain (the difference in body weight from 1 to 12mo) in both females (E) and males (F). **(G-I)** Significant effect of running on the change in lean mass (G), fat mass (H), and fat percentage (I) between six and eleven months, with an overall reduction in running mice compared to sedentary mice across all *APOE* genotypes in females. **(J-L)** Running had a significant reduction on the change in lean mass (J) and fat mass (K) between six and eleven months, but no change in fat percentage (L) in male mice. Data presented as mean ± SEM, two-way ANOVA performed for *APOE* genotype (significant marked above ‘Sed’ column, indicating an effect of *APOE* genotype), Running (significance marked above ‘Run’ column, indicating an effect of running), and the interaction between *APOE* genotype:Running (significance marked to the right of the graph). Bonferroni’s multiple comparisons performed for within genotype running effects (significance marked in smaller stars directly to the right of the run column, within graph limits, in the color of the genotype). *P < 0.05, **P < 0.01, ***P < 0.001, ****P < 0.0001.

Overall, running mice had a lower fat composition compared to sedentary mice for both sexes at 6 and 11mo (**Supp Fig 4,5**). In females, only *APOE^ε4/ε4^* mice showed a significant attenuation of fat mass and fat percentage in running compared to sedentary mice (**Fig 2G-I, Supp Fig 3E-G**). There were no *APOE* genotype differences in male mice, however there was an effect of running on lean and fat mass. This effect was most pronounced in *APOE^ε3/ε4^* male mice, with running attenuating lean and fat mass (**Fig 2J-L, Supp Fig 3H-J**). Running attenuated the age-related increase in lean and fat mass across all genotypes and sexes. However, there was a pronounced reduction of age-related fat mass accumulation in female *APOE^ε4/ε4^* running mice. Also, male *APOE^ε3/ε4^* running mice showed the greatest reduction in age-related lean and fat mass accumulation.

No effect of running or *APOE* genotype was determined for total cholesterol or HDL concentration at 12mo (**Supp Fig 6**). There was a significant sex-specific effect of *APOE* genotype on LDL concentration in the plasma. In running females, LDL concentrations decreased in an *APOE^ε4^* dose-dependent manner (**Supp Fig 6H**). Conversely, in running males, LDL concentrations were significantly lower than sedentary mice (**Supp Fig 6K**). Cholesterol composition in running mice did not correlate with running distance for both sexes (**Supp Fig 6C,F,I,L,O,R**).

Collectively, these data demonstrate that weight, body composition and cholesterol levels, commonly used markers of healthy aging, are significantly affected by voluntary running, but the effects are dependent upon both sex and *APOE* genotype.

### Running affects activities of daily living in an *APOE* genotype- and sex-specific manner

In humans, prior to more severe cognitive decline in ADRDs, activities of daily living (i.e., sleep, general movement, feeding) are often disrupted[50–52]. To evaluate activities of daily living in mice, we measured feeding and walking behavior (pedometers) across four dark cycles (active/awake period), and three light cycles (inactive/sleep period) at 11 mos. Feeding behavior revealed significant changes in the dark cycle, but not in the light cycle for both sexes. In females, running mice consumed more food than sedentary mice during the dark cycle across all *APOE* genotypes (**Fig 3A-B, Supp Fig 7**). However, in males, there was an interaction between *APOE* genotype and running during the dark cycle. In sedentary mice, *APOE^ε3/ε4^* ate more than *APOE^ε3/ε3^* and *APOE^ε4/ε4^*. This pattern was not apparent in running mice, suggesting running mitigates the *APOE* genotype differences observed in sedentary mice (**Fig 3D, Supp Fig 7**).

**Figure 3:**
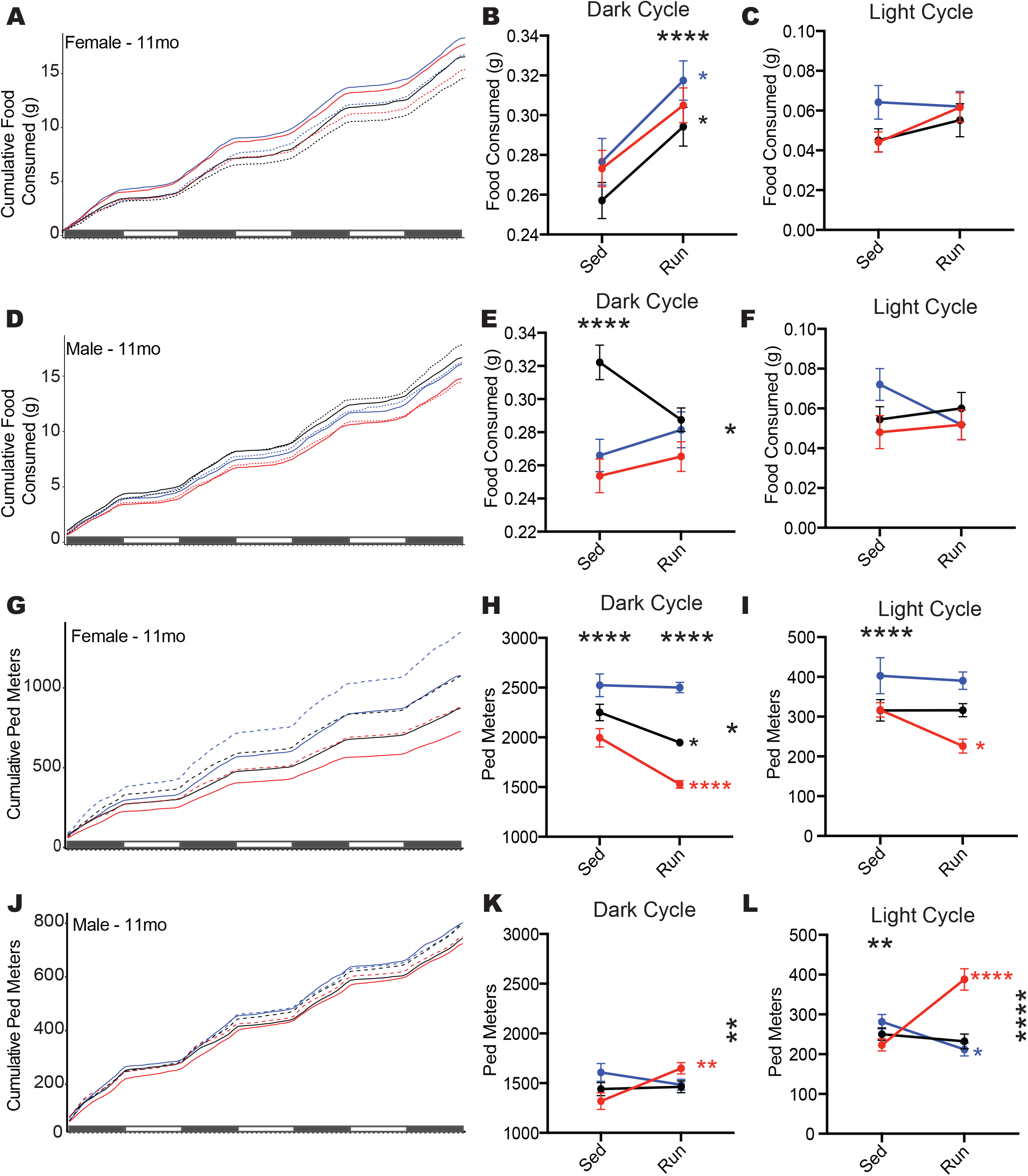
Activities of daily living are influenced by *APOE* genotype and running. **(A,D)** Cumulative food consumed per gram for female (A) and male (D) mice across four dark cycles and three light cycles. **(B-C)** Running significantly increased food consumed during the dark cycle (B), but not the light cycle (C) in female mice. **(E-F)** *APOE* genotype and *APOE* genotype:running interaction affected food consumption in males during the dark cycle (E), but no effect was seen during the light cycle (F). **(G,J)** General movement (cumulative ped meters) for female (G) and male (J) mice across four dark cycles and three light cycles. **(H-I)** *APOE* genotype, Running, and *APOE* genotype:Running interaction all significantly affected ped meters during the dark cycle (H) with running decreasing ped meters differently across the genotypes. Only *APOE* genotype was significant during light cycle (I) in female mice. **(K-L)** *APOE* genotye:Running interaction significantly affected ped meters in males, with *APOE^ε4/ε4^* showing increased ped meters during the dark cycle (K), as well as the light cycle (L). There wads also an *APOE* genotype effect (L). Data presented as mean ± SEM, two-way ANOVA performed for *APOE* genotype (significant marked above ‘Sed’ column, indicating an effect of APOE genotype), Running (significance marked above ‘Run’ column, indicating an effect of running), and *APOE* genotype:Running interaction (significance marked in to the right of the graph). Bonferroni’s multiple comparisons performed for within genotype running effects (significance marked in smaller stars directly to the right of the run column, within graph limits, in the color of the genotype). *P < 0.05, **P < 0.01, ***P < 0.001, ****P < 0.0001.

We next determined whether movement in the home cage was affected by *APOE* genotype and/or running by measuring walking (Ped meters) (**see Methods**). Female sedentary mice showed an *APOE^ε4^* dose-dependent increase in Ped meters that was attenuated by running (**Fig 3G-H, Supp Fig 7**). During the light cycle only *APOE^ε4/ε4^* females showed a significant reduction in Ped meters compared to their sedentary counterparts (**Fig 3I, Supp Fig 7**). In male mice, both *APOE* genotype and running interacted to alter Ped meters during both the dark and light cycle, however running more strikingly increased cumulative Ped meters of *APOE^ε4/ε4^* mice compared to the other *APOE* genotypes (**Fig 3K-L, Supp Fig 7**).

These results show that *APOE* genotype modulates the effects of running on natural home cage behaviors such as feeding and general movement, considered equivalent to activities of daily living in humans.

### *APOE* genotype affects running-dependent increase in energy expenditure during the dark cycle

Previous studies in humans demonstrated *APOE* genotype affects metabolism on a cellular, regional, and organismal level[53, 54]. To determine whether running and *APOE* genotype affect metabolic processes, energy expenditure (kcal/hr) was measured at 11mo. In female sedentary mice, energy expenditure showed an *APOE^ε4^* dose-dependent increase during the dark cycle. In general, running resulted in significantly higher energy expenditure in the dark cycle in male and female mice. However, this effect was not observed in male *APOE^ε3/ε4^* mice (**Fig 4A-C, Supp Fig 8**). This suggests the *APOE^ε3/ε4^* genotype attenuates the effects of running in a sexually dimorphic manner. During the light cycle, all genotypes showed decreased energy expenditure with running; except for female *APOE^ε3/ε3^* mice that showed an increase (**Fig 4D-F, Supp Fig 8**). Subtle but significant changes in substrate usage (based on respiratory quotient, RQ, **see methods**) were also determined across groups in both the light and dark cycle (**Fig 4G-L, Supp Fig 8**). Significant changes were small, however may worsen with more advancing age (**Fig 4H-L, Supp Fig 8**). These results highlight that *APOE* genotype and running affect energy expenditure, however changes in starting energy substrate usage were minute (**Fig 4A-F**).

**Figure 4:**
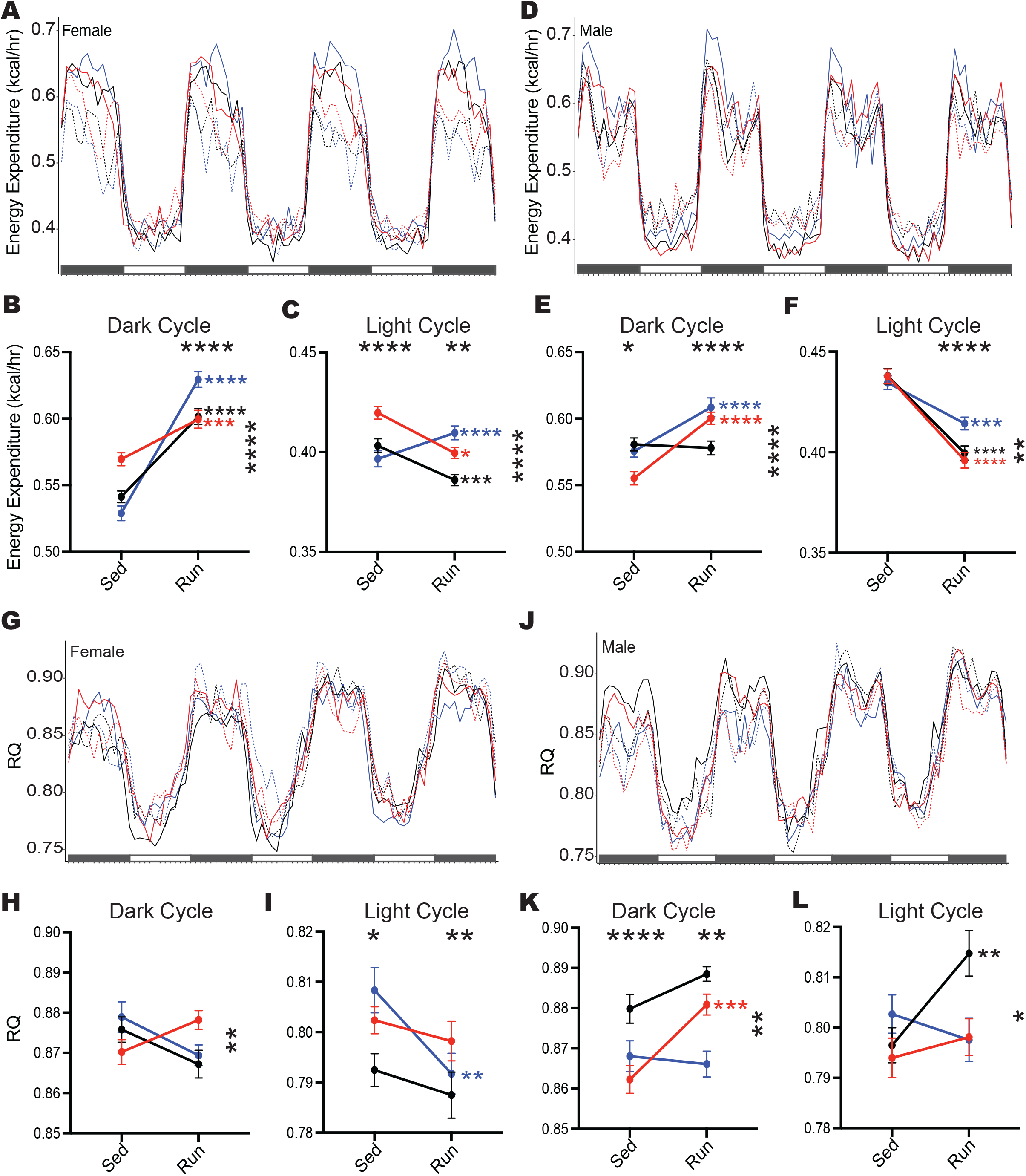
Running influences energy expenditure differently between females and male *APOE* mice. **(A-C**) Energy expenditure (kcal/hr) across four dark cycles and three light cycles for female mice. Energy expenditure (kcal/hr) significantly affected by *APOE* gentoype:Running, and Running during the dark cycle (B), with an increase in energy expenditure in running mice. *APOE* genotype:Running, *APOE* genotype, and Running all influenced light cycle energy expenditure (C), with Apoeε3/3 increasing with running while *APOE^ε3/ε3^* and *APOE^ε4/ε4^* decreased energy expenditure with running. **(D-F)** Energy expenditure (kcal/hr) across four dark cycles and three light cycles for male mice. (E) *APOE* genotype:Running, *APOE* genotype, and Running all affected dark cycle energy expenditure in male mice. (F) *APOE* genotype:Running and Running showed an overall decrease in energy expenditure in running male mice. **(G-I)** Respiratory Quotient (RQ) across four dark cycles and three light cycles for female mice (G). *APOE* genotype:Running significantly affected RQ during the dark cycle for female mice (H). *APOE* genotype and Running significantly affected RQ during the light cycle for female mice (I). **(J-L)** Respiratory Quotient (RQ) across four dark cycles and three light cycles for male mice (J). *APOE* genotype:Running, *APOE* genotype, and Running all significantly affected RQ during the dark cycle in male mice (K). *APOE* genotype:Running significantly affected RQ during the light cycle (L). Data presented as mean ± SEM, two-way ANOVA performed for *APOE* genotype (significant marked above ‘Sed’ column, indicating an effect of *APOE* genotype), Running (significance marked above ‘Run’ column, indicating and effect of running), and the interaction between *APOE* genotype:Running (significance marked to the right of the graph). Bonferroni’s multiple comparisons performed for within genotype running effects (significance marked in smaller stars directly to the right of the run column, within graph limits, in the color of the genotype). *P < 0.05, **P < 0.01, ***P < 0.001, ****P < 0.0001.

### *APOE* genotype causes subtle sex-specific changes to the effects of running on the aging brain

Unbiased transcriptional profiling was utilized to capture molecular effects across *APOE* genotype and activities (12 groups per brain region, **Fig. 5A, See Methods**). Principal component analysis (PCA) revealed brain region (PC1, 65%) and sex (PC2, 20%) were the primary drivers of variance, suggesting *APOE* genotype and running are not exerting strong effects (**Fig 5B**). Therefore, to determine subtle effects of *APOE* genotype and running, linear modeling was used for male or female cortex or hippocampus samples separately (4 linear models in total) (**Fig 5C**). Supporting the PCA data, linear modeling revealed fewer than 200 significant genes in females for the cortex and hippocampus, and fewer than 800 genes in males. These numbers align with published data from human studies but are somewhat fewer than previous mouse studies (**Supp Fig 12**). Several significant genes including *Ephx1* (main effect: *APOE^ε3/ε4^*), *Ctsf* (main effect: *APOE^ε4/ε4^*), *C3* (interaction *APOE^ε3/ε4^*:Run) and *Cav3* (interaction *APOE^ε4/ε4^*:Run) are known to function in lipid homeostasis, neuroinflammation and membrane integrity, key processes implicated in ADRD risk (**Fig 5H**).

**Figure 5:**
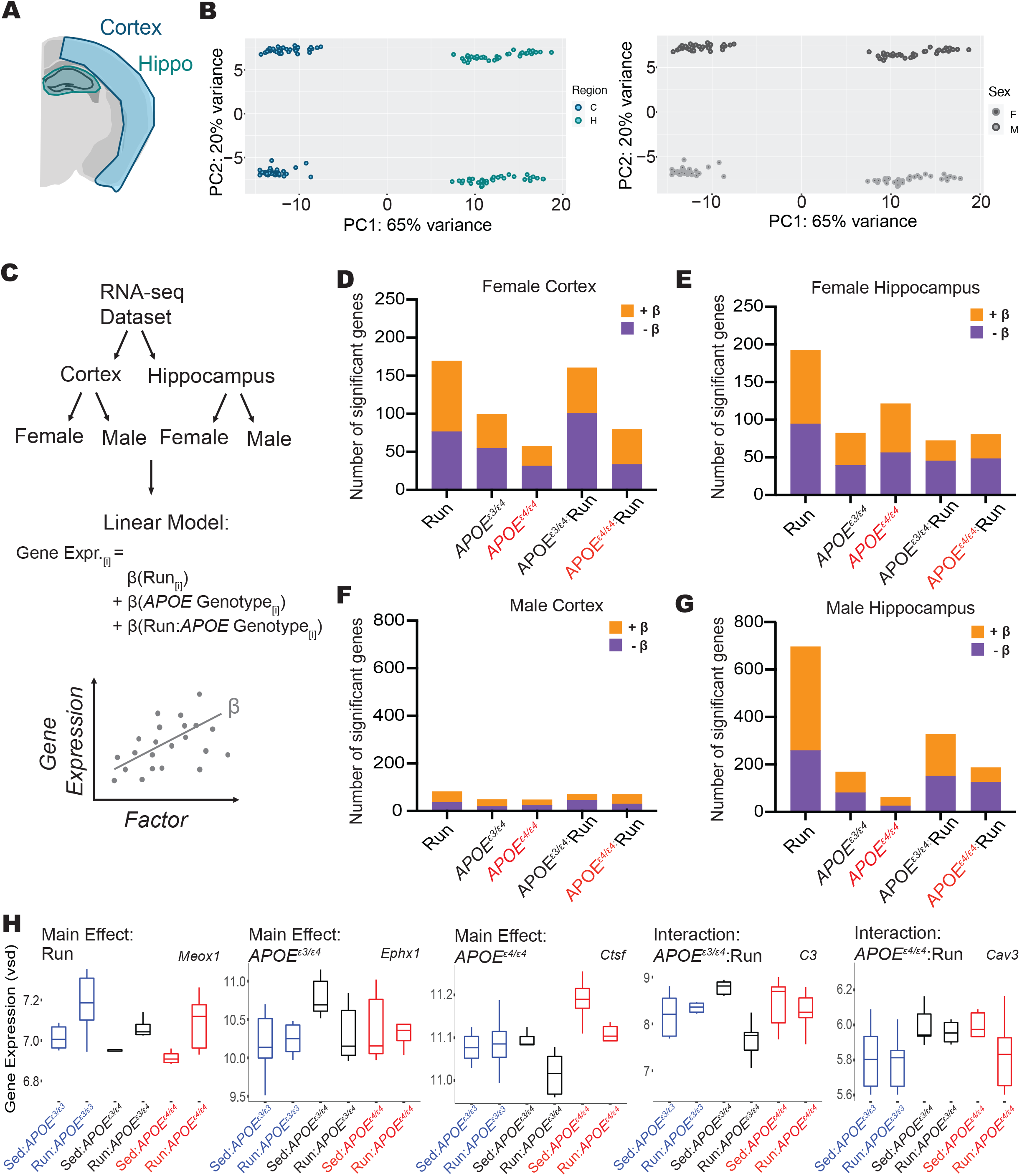
Transcriptional profiling reveals subtle changes due to *APOE* genotype and running in the cortex and hippocampus. **(A)** Diagram of the cortical and hippocampal regions of the brain taken for transcriptional profiling. **(B)** PCA revealed clear separations between brain regions (cortex and hippocampus, 65% variance explained), as well as by sex (female and male, 20% of variance explained). **(C)** Schematic of the computational analysis approach; first RNA-seq was separated by brain region, next separated again by sex. Four linear models were run to examine gene expression as it varies with Running, *APOE* genotype, and the interaction between *APOE* genotype:Running. β-value is the association of the gene with the factor tested – positive β-value indicates a positive correlation, negative β-value indicates a negative correlation. **(D-G)** Number of significant genes (FDR corrected) for female cortex (D), female hippocampus (E), male cortex (F), and male hippocampus (G). **(H)** Example of a significant gene for each of the main effects and interactive effects: Meox1 (Hippocampus, Male), Ephx1 (Hippocampus, Male), Ctsf (Hippocampus, Female), C3 (Hippocampus, Male), Cav3 (Cortex, Male).

For the female cortex, GO terms showed positive NES for the main effects (running, *APOE^ε3/ε4^*, and *APOE^ε4/ε4^*), but negative NES for the interactive terms (*APOE^ε3/ε4^*:Run, *APOE^ε4/ε4^*:Run) for vascular and synaptic/neuronal functions (**Fig 6B-C**). Also, interestingly, in males, there were few significantly enriched GO terms for *APOE^ε3/ε4^*, suggesting in males, but not females, the *APOE^ε3/ε4^* genotype exerts little to no effect compared to the *APOE^ε3/ε3^* genotype on genes associated with vascular integrity-related processes (**Fig 6D**). Transcriptional profiling supplemented our peripheral findings, determining that running and *APOE* genotype interact in sex-specific ways to influence mechanisms involved in dementia-relevant biological processes.

**Figure 6:**
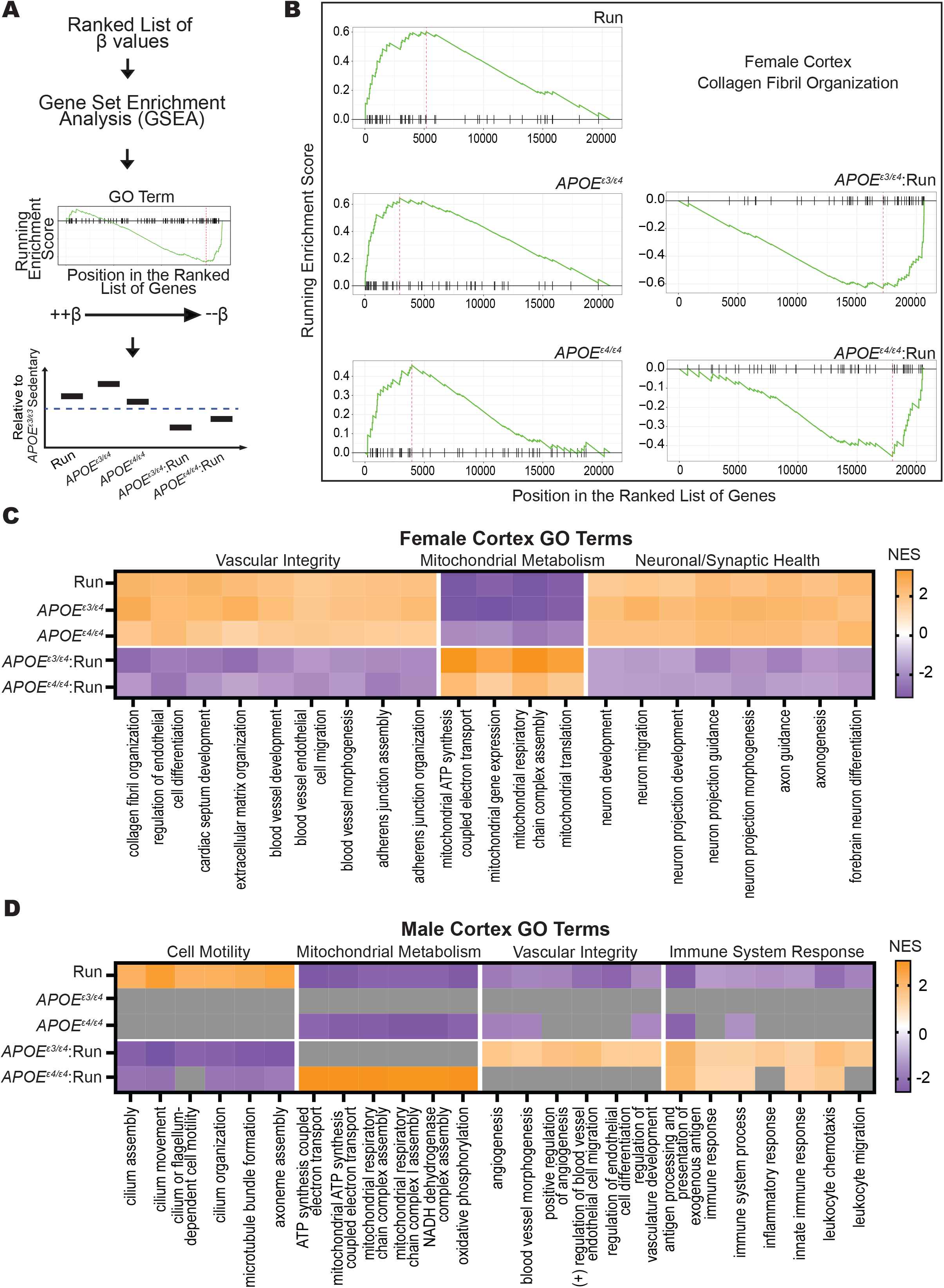
GSEA predicts *APOE* genotype and running interact to mitigate main effects. **(A)** Schematic of computational approach; β-values from linear models were passed through GSEA for gene ranking, GSEA plots were used to visualize results and main effects of running, *APOE* genotype, and *APOE* genotype:Running were interpreted per GO term. **(B)** GSEA plots for ‘Collagen Fibril Organization’ in the female cortex. Main effects of Running and *APOE^ε3/ε4^*, and *APOE^ε4/ε4^* all show positive Enrichment scores, while the interactions, *APOE^ε3/ε4^*:Run and *APOE^ε4/ε4^*:Run reveal negative Enrichment Scores. **(C)** In the female cortex data GO terms that fit the pattern shown in (B), colored by Normalized Enrichment Score (NES), are represented specifically vascular integrity, mitochondrial metabolism, and synaptic/neuronal health. **(D)** The pattern observed in male cortex was different to that seen in female cortex (B,C) with *APOE^ε3/ε4^* appearing more similar to *APOE^ε3/ε3^* (indicated by gray boxes) for enrichment terms grouped as cell motility, mitochondrial metabolism, vascular integrity, and immune system response.

## 4. Discussion

Exercise is generally considered to have beneficial effects, but our results show *APOE* genotype impacts the effects of running. Significant interactions between *APOE* genotype and running were observed across body weight, body composition, activities of daily living, systemic metabolism, and cortical and hippocampal gene expression. Male and female mice were evaluated separately as ADRD risk varies between the sexes, with higher risk in women compared to men[55, 56]. Sex is typically used as a covariate in human studies, but our data show that *APOE* genotype and sex interact across multiple domains. Additionally, there is a lack of consideration that odds ratios are sex-specific when assessing clinical trials, obfuscating the effects of sex. Our data suggest *APOE* genotype for each sex should be considered for studies assessing exercise interventions to reduce risk for dementia.

While the brain has been shown to be plastic throughout adulthood, environmental influences can exert a greater effect on a younger brain compared to an older brain[19, 23, 35, 57–60], prompting us to study the effects of *APOE* genotype and running from early age to midlife. We assessed 12mo mice to understand the effect of *APOE* and running up until midlife, likening our findings to prodromal studies in humans[61, 62]. *APOE* genotype-specific effects may also be apparent at older ages so studying later timepoints in the mouse, even beginning running at midlife would be informative. This would relate more closely with human clinical trials that conduct studies on older, affected human populations (i.e., nursing home/hospice patients)[63–65].

Transcriptomic approaches have revolutionized our understanding of ADRDs and has therefore become a hypothesis generating tool for identifying the molecular pathways impacted by genetic and environmental risk factors. Therefore, we used transcriptional profiling to identify interactions between *APOE* genotype and running. Our data revealed a reversal of NES direction from the main effects and the interaction of *APOE* genotype and running. This was unexpected, as we saw similar patterns of positive (or negative) enrichment for 1) running compared to sedentary, 2) *APOE^ε3/ε4^* compared to *APOE^ε3/ε3^* and 3) *APOE^ε4/ε4^* compared to *APOE^ε3/ε3^*. These results contradict the assumption that running would have the opposite effect on the brain as *APOE^ε4^*, particularly when considering each of these terms collectively (vascular, immune, mitochondrial, neuronal/synaptic). We propose that there is a possibility for overcompensation for the *APOE^ε4^* allele. While evidence shows *APOE^ε4^* causes gains and losses of APOE function across many processes, it is unknown whether there is a preemptive response that has not been considered. Further, the *APOE^ε3/ε4^* genotype may be responding to early aging phenotypes different than *APOE^ε4/ε4^* genotype. Precise experimentation on this phenomenon is needed in both mice and humans to better understand which *APOE^ε4^*-specific pathways are mitigated by running. Lastly, while these models are key for interpretation of APOE biology, other important pathological interactions (e.g., amyloid or TAU) are not present in this study. Future studies are necessary to interrogate the interaction between APOE, exercise, and hallmark ADRD pathologies in order to provide further translatable outcomes.

Advancements in RNA-sequencing have made it cheaper and faster to sequence the brains of ADRD patients (ROSMAP, MAYO, ADSP). Recently, research programs have explored whether *APOE* influences the human cerebral transcriptome. In three largescale AMP-AD studies, reports included few to no changes in multiple brain regions in *APOE^ε4^*+ cases (ROSMAP: syn8456629, MAYO: syn8466812, MSBB: syn8484987)[66–68] (**Supp Fig 12**). The ROSMAP dataset analysis showed no differences due to *APOE^ε4^* status across the dorsolateral prefrontal cortex region[66]. The MAYO dataset showed a significant differential expression (DE) of only 173 genes between *APOE^ε3/ε4^* and *APOE^ε3/ε3^*, and a significant DE of only 88 genes between *APOE^ε4/ε4^* and *APOE^ε3/ε3^* in the temporal cortex[67]. The Mount Sinai Brain Bank (MSBB) reported fewer than 5 genes DE between all *APOE* genotype comparisons in the frontal pole region, parahippocampal gyrus region, frontal superior temporal gyrus region, and inferior frontal gyrus region[68]. Our mouse data aligns more closely with these human studies, possibly due to litter-matched mice, and further analyses using GSEA showed subtle changes that escaped detection through traditional DE analysis. Moving forward, our data show the importance of including heterozygous genotypes (e.g., *APOE*^ε3/ε4^) and varying degrees of chronic voluntary exercise (e.g., low, medium, high) in mouse studies to improve the alignment to ADRD in human studies.

The *APOE^ε4^* allele emerged as our early hominin ancestors adapted to changes in habitat and food availability to include more aerobic exercise such as running[69]. The *APOE^ε4^* allele was beneficial for storage of fats, increasing cholesterol. While the *APOE^ε4^* conferred longer lifespan 200,000 years ago, the diet and exercise of an individual was drastically different[69]. Currently, western culture sees some of the highest rates of ADRD, due to the interaction between *APOE^ε4^* and our environment, and as we show, running. This work supports that *APOE^ε4^* interacts with running in a genotype- and sex-specific manner, influencing ADRD risks in the periphery and brain.

## Author contributions

KEF and GRH conceived and designed this project. KEF, CAD, AAH performed mouse experiments. KEF performed IF, experimental analysis, and bioinformatic analysis. KEF and GRH consulted for statistical approach and analysis. KEF and GRH wrote and prepared this manuscript. All authors read and approved the final manuscript.

## Acknowledgements

Research reported in this publication was partially supported by the Jackson Laboratory’s Genetic Engineering Technologies Scientific Service. The authors wish to thank Todd Hoffert from Clinical Assessment Services for blood chemistry, Heidi Munger and the Genome Technologies group for RNA-sequencing. We would also like to thank Dr. Gregory Carter and Dr. Christoph Pruess for their continued advice on computational analysis and support for these projects.

## Declarations

The authors are supported by funding from the National Institutes of Health, National Institute on Aging U54 AG054345.

## Ethics Approval and Consent to Participate

No human subjects or data was used in this study. All experiments involving mice were approved by the Animal Care and Use Committee at The Jackson Laboratory in accordance with guidelines set out in The Eighth Edition of the Guide for the Care and Use of Laboratory Animals. All euthanasia used methods were approved by the American Veterinary Medical Association.

## Availability of Data and Materials

All data is being uploaded to the AD Knowledge Portal. ID numbers will be provided once the process is complete.

