## Supplemental Data 1 for "APOE^ε4^ and exercise interact to influence systemic and cerebral risk factors for dementia"

**Supp Fig. 1**

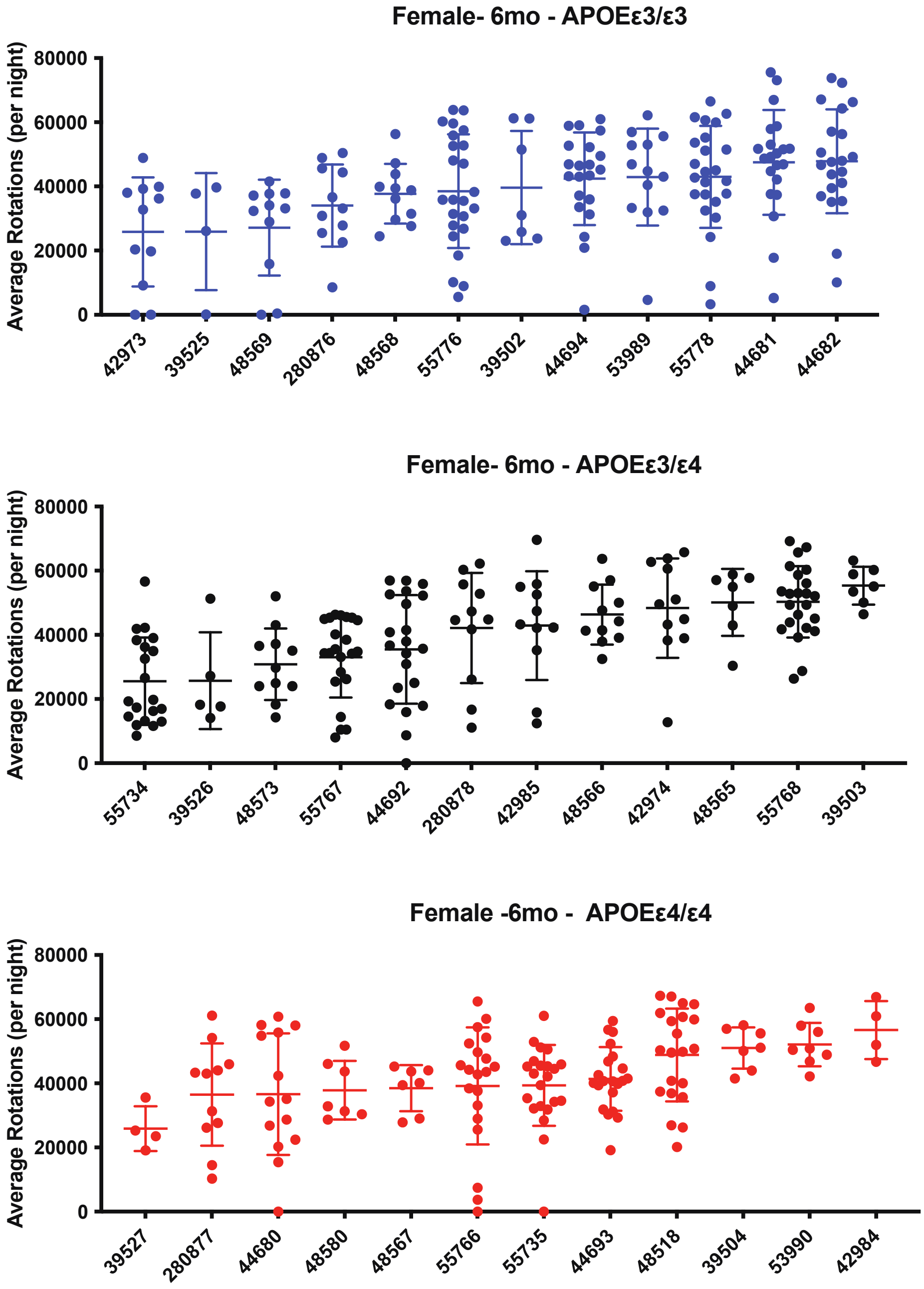


**Supp Fig. 1**
Average Rotations (per night) for each female mouse measured across APOE genotypes at 6mo. Each dot represents a night. Variation was expected within genotypes. Data presented as mean ± SD.

**Supp Fig. 2**

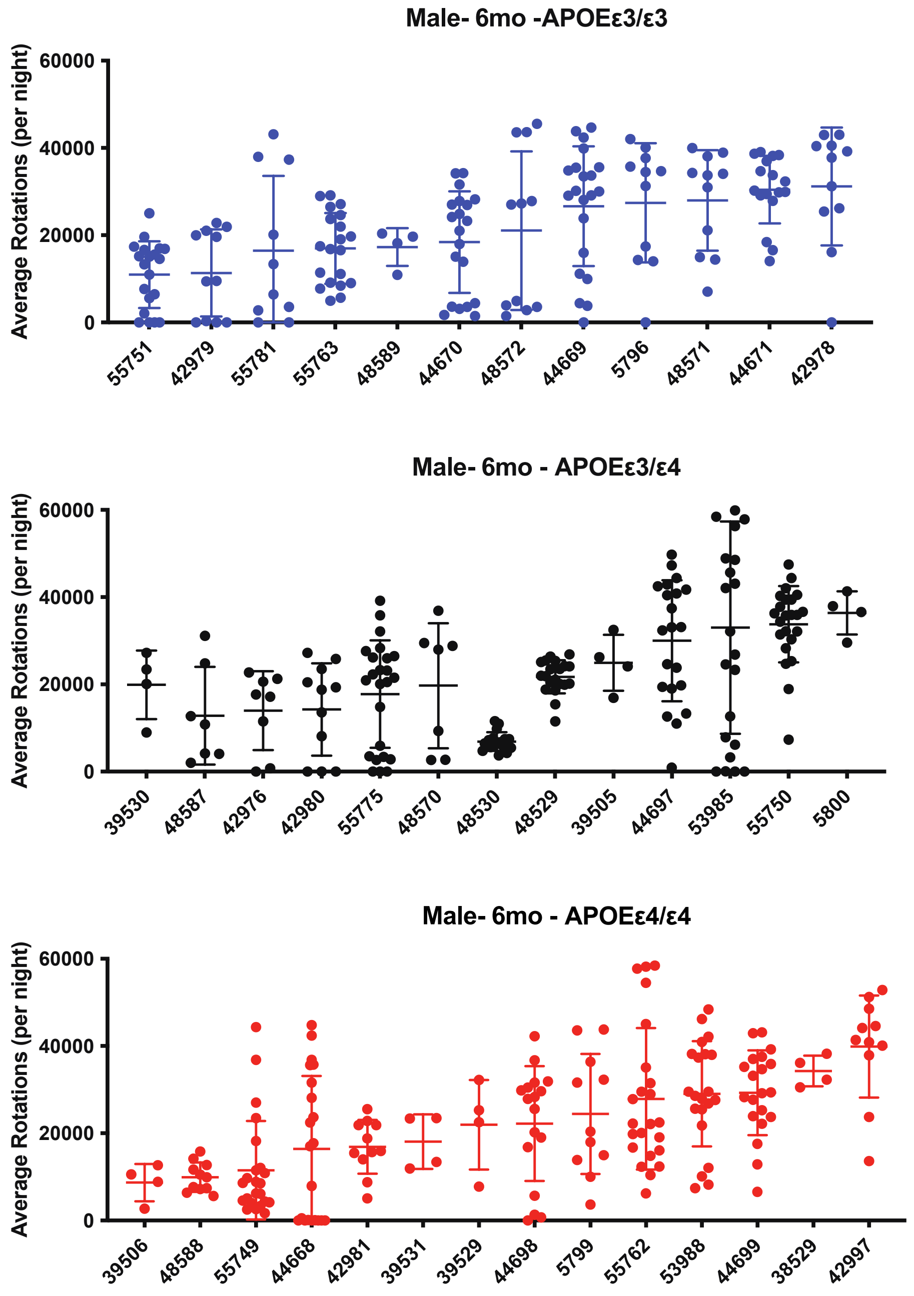


**Supp Fig. 2**
Average Rotations (per night) for each male mouse measured across APOE genotypes at 6mo. Each dot represents a night. Variation was expected within genotypes. Data presented as mean ± SD.

**Supp Fig. 3**

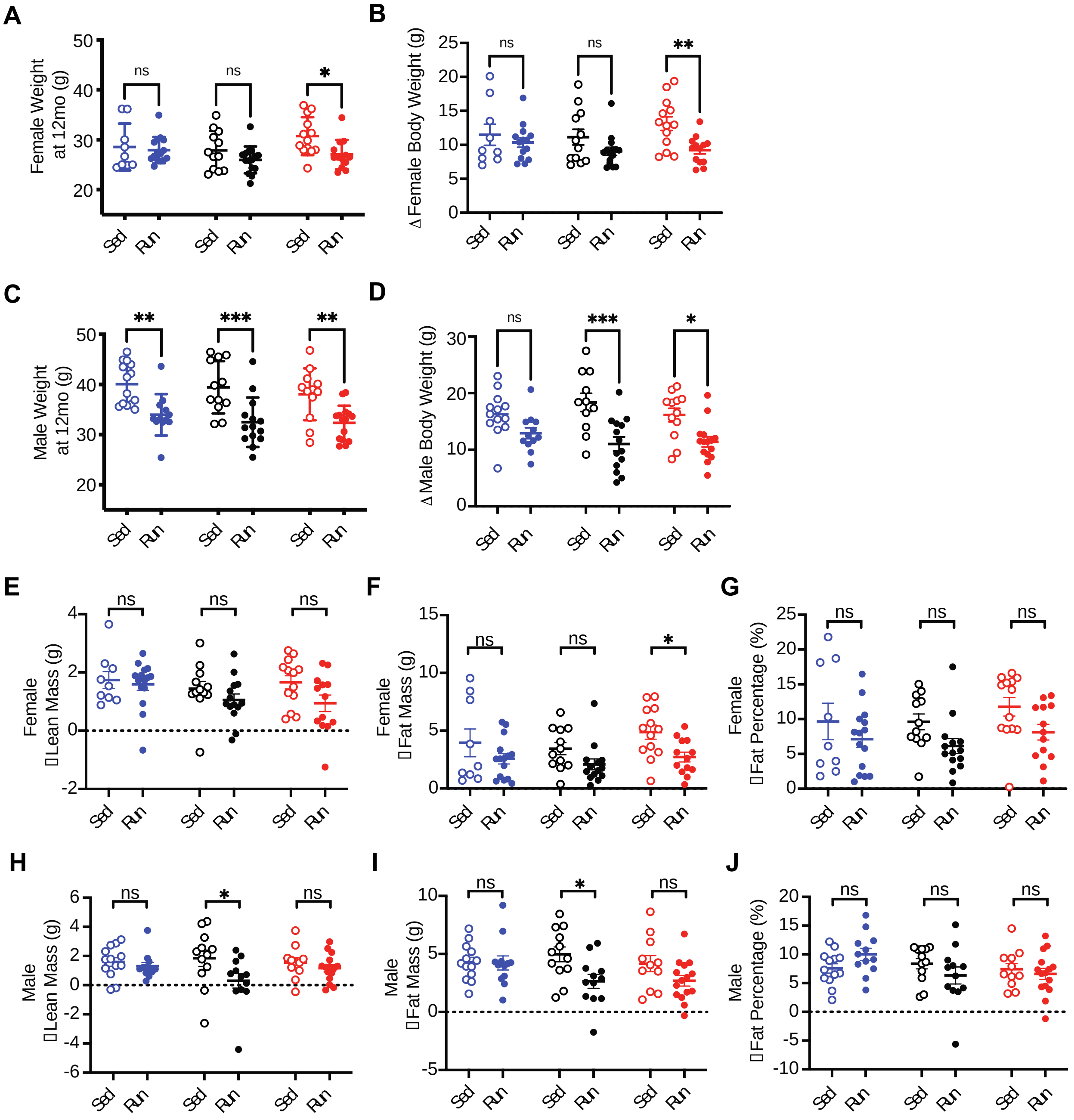


**Supp Fig. 3**
**(A)** Bonferonni posthoc results for female weight (g) at 12mo. **(B)** Bonferonni posthoc results for the change in female weight (g) between 1 and 12mo. **(C)** Bonferonni posthoc results for male weight (g) at 12mo. **(D)** Bonferonni posthoc results for the change in male weight (g) between 1 and 12mo. **(E-G)** Bonferonni posthoc results for the change in female lean mass (E), fat mass (F), and fat percentage (G) from 6mo to 11mo. **(H-J)** Bonferonni posthoc results for the change in male lean mass (H), fat mass (I), and fat percentage (J) from 6mo to 11mo. Data presented as mean ± SEM.

#### Supp Fig. 4

6 months

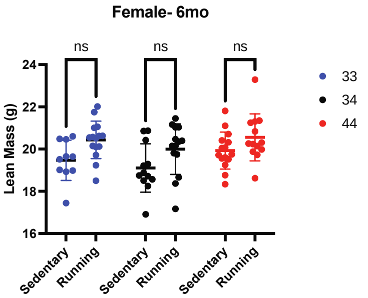


| Source of Variation | % of total variation | P value | P value summary |
|---|---|---|---|
| Interaction | 0.4454 | 0.8171 | ns |
| Activity | 13.61 | 0.0008 | *** |
| Genotype | 6.625 | 0.0553 | ns |

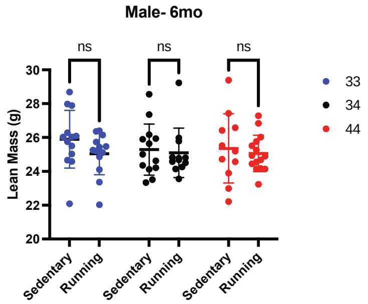


| Source of Variation | % of total variation | P value | P value summary |
|---|---|---|---|
| Interaction | 0.9685 | 0.7001 | ns |
| Activity | 2.262 | 0.1999 | ns |
| Genotype | 0.7187 | 0.7673 | ns |

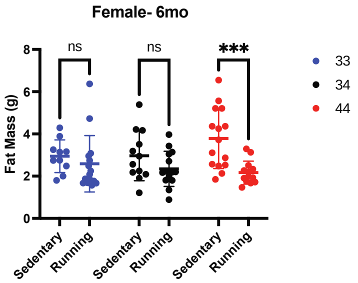


| Source of Variation | % of total variation | P value | P value summary |
|---|---|---|---|
| Interaction | 5.256 | 0.0938 | ns |
| Activity | 13.14 | 0.0008 | *** |
| Genotype | 1.263 | 0.5584 | ns |

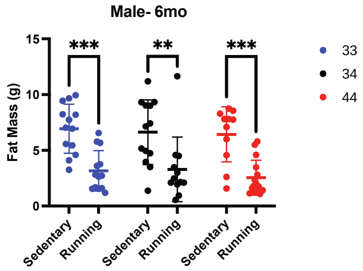


| Source of Variation | % of total variation | P value | P value summary |
|---|---|---|---|
| Interaction | 0.1634 | 0.9063 | ns |
| Activity | 39.05 | <0.0001 | **** |
| Genotype | 0.6939 | 0.6596 | ns |

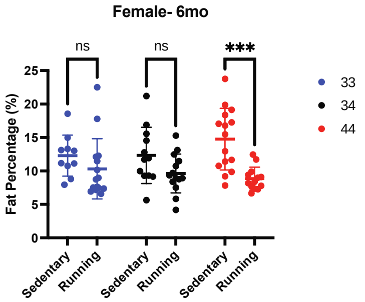


| Source of Variation | % of total variation | P value | P value summary |
|---|---|---|---|
| Interaction | 4.356 | 0.1276 | ns |
| Activity | 18.01 | <0.0001 | **** |
| Genotype | 0.6972 | 0.7136 | ns |

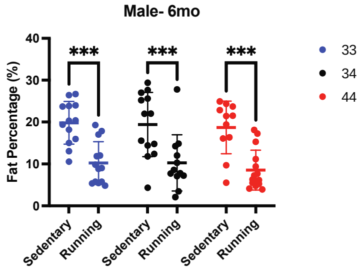


| Source of Variation | % of total variation | P value | P value summary |
|---|---|---|---|
| Interaction | 0.07817 | 0.9531 | ns |
| Activity | 40.28 | <0.0001 | **** |
| Genotype | 0.6589 | 0.6685 | ns |

**Supp Fig. 4**

Bonferroni posthoc results for body composition data for females (left column) and males (right column) at 6mo. Table below figures are results from two-way ANOVA, significance on graphs is from Bonferonni posthoc results. Data presented as mean ± SD. *P < 0.05, **P < 0.01, ***P < 0.001, ****P < 0.0001.

#### Supp Fig. 5

11 months

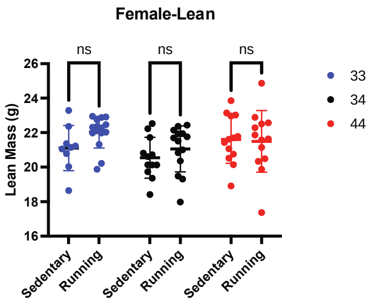


| Source of Variation | % of total variation | P value | P value summary |
|---|---|---|---|
| Interaction | 2.348 | 0.3966 | ns |
| Activity | 2.529 | 0.1598 | ns |
| Genotype | 6.771 | 0.0740 | ns |

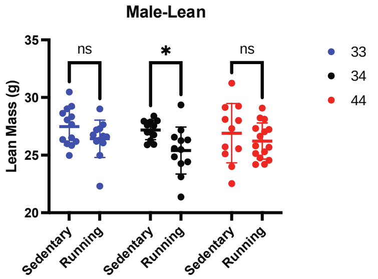


| Source of Variation | % of total variation | P value | P value summary |
|---|---|---|---|
| Interaction | 1.531 | 0.5437 | ns |
| Activity | 10.33 | 0.0053 | ** |
| Genotype | 2.160 | 0.4246 | ns |

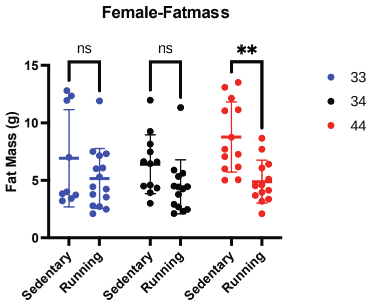


| Source of Variation | % of total variation | P value | P value summary |
|---|---|---|---|
| Interaction | 2.432 | 0.3316 | ns |
| Activity | 16.89 | 0.0002 | *** |
| Genotype | 3.635 | 0.1944 | ns |

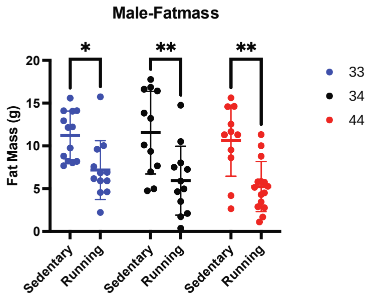


| Source of Variation | % of total variation | P value | P value summary |
|---|---|---|---|
| Interaction | 0.6013 | 0.7277 | ns |
| Activity | 31.66 | <0.0001 | **** |
| Genotype | 1.426 | 0.4728 | ns |

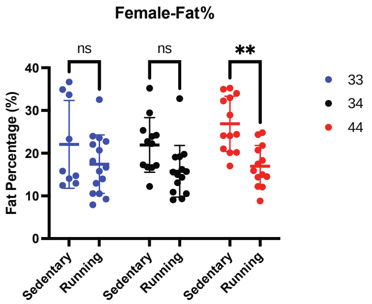


| Source of Variation | % of total variation | P value | P value summary |
|---|---|---|---|
| Interaction | 2.081 | 0.3756 | ns |
| Activity | 20.16 | <0.0001 | **** |
| Genotype | 2.927 | 0.2542 | ns |

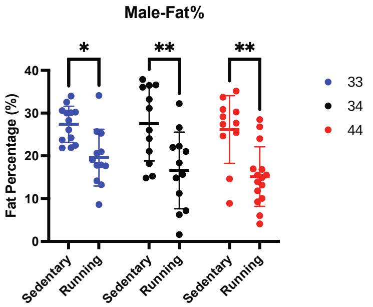


| Source of Variation | % of total variation | P value | P value summary |
|---|---|---|---|
| Interaction | 0.7305 | 0.6777 | ns |
| Activity | 31.61 | <0.0001 | **** |
| Genotype | 1.745 | 0.3975 | ns |

**Supp Fig. 5**
Bonferroni posthoc results for body composition data for females (left column) and males (right column) at 12mo. Table below figures are results from two-way ANOVA, significance on graphs is from Bonferonni posthoc results. Data presented as mean ± SD. *$P < 0.05$, **$P < 0.01$, ***$P < 0.001$, ****$P < 0.0001$.

#### Supp Fig. 6

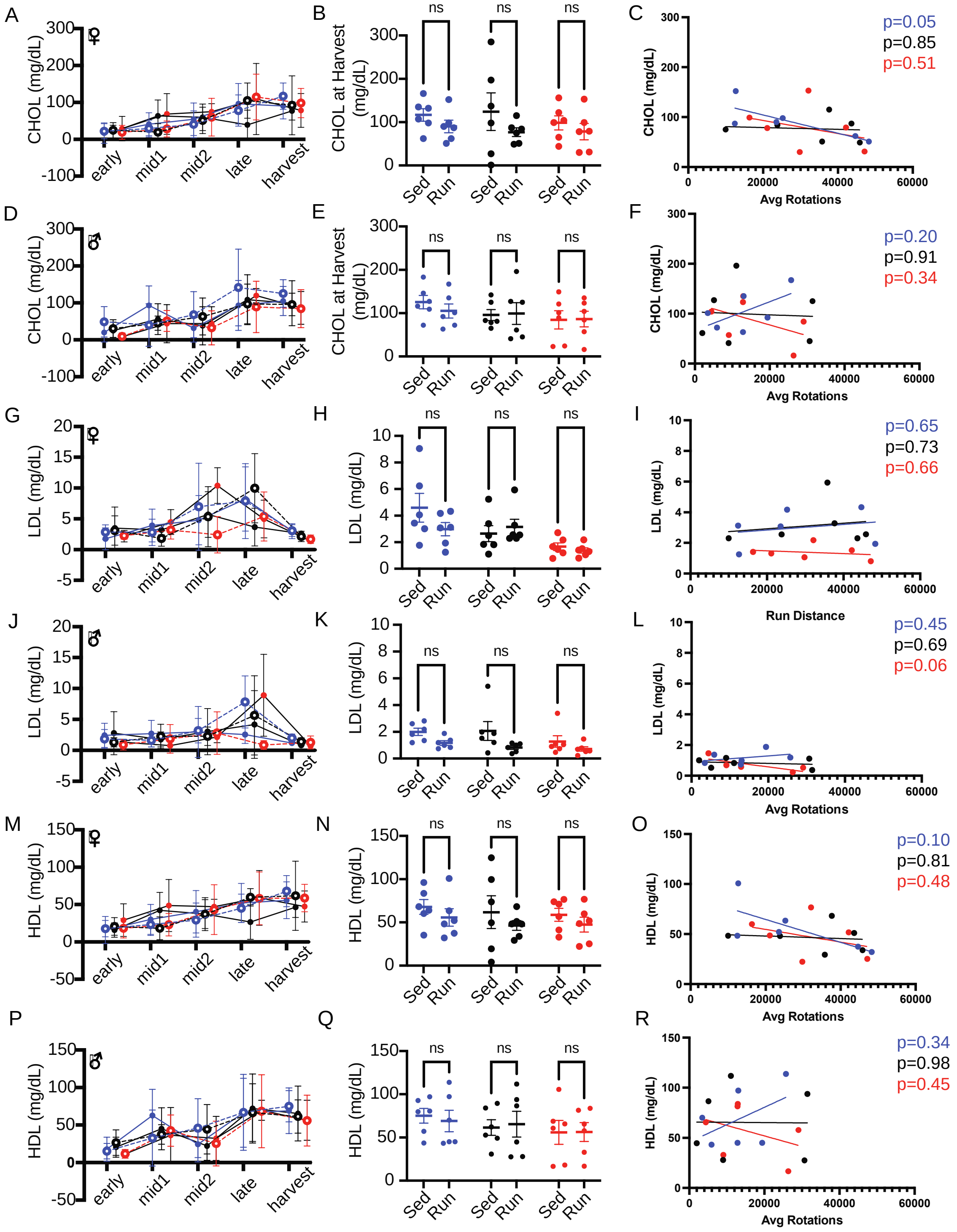


**Supp Fig. 6**

**(A-C)** Female total cholesterol (CHOL, mg/dL) measured across the experiment (A), at harvest (B), and correlation with running distance at 11mo (C). **(D-F)** Male total cholesterol (CHOL, mg/dL) measured across the experiment (D), at harvest (E), and correlation with running distance at 11mo (F). **(G-I)** Female LDL (mg/dL) measured across the experiment (G), at harvest (H), and correlation with running distance at 11mo (I). **(J-L)** Male LDL (mg/dL) measured across the experiment (J), at harvest (K), and correlation with running distance at 11mo (L). **(M-O)** Female HDL (mg/dL) measured across the experiment (M), at harvest (N), and correlation with running distance at 11mo (O). **(P-R)** Male HDL (mg/dL) measured across the experiment (P), at harvest (Q), and correlation with running distance at 11mo (R). Five timepoints: early (0-2 months), mid1 (3-5 months), mid2 (6-8 months), late (9-11 months), and harvest (12 months). Data presented as mean ± SD (A,D,G,J,M,P). Data presented as mean ± SEM (B,E,H,K,N,Q)

**Supp Fig. 7**

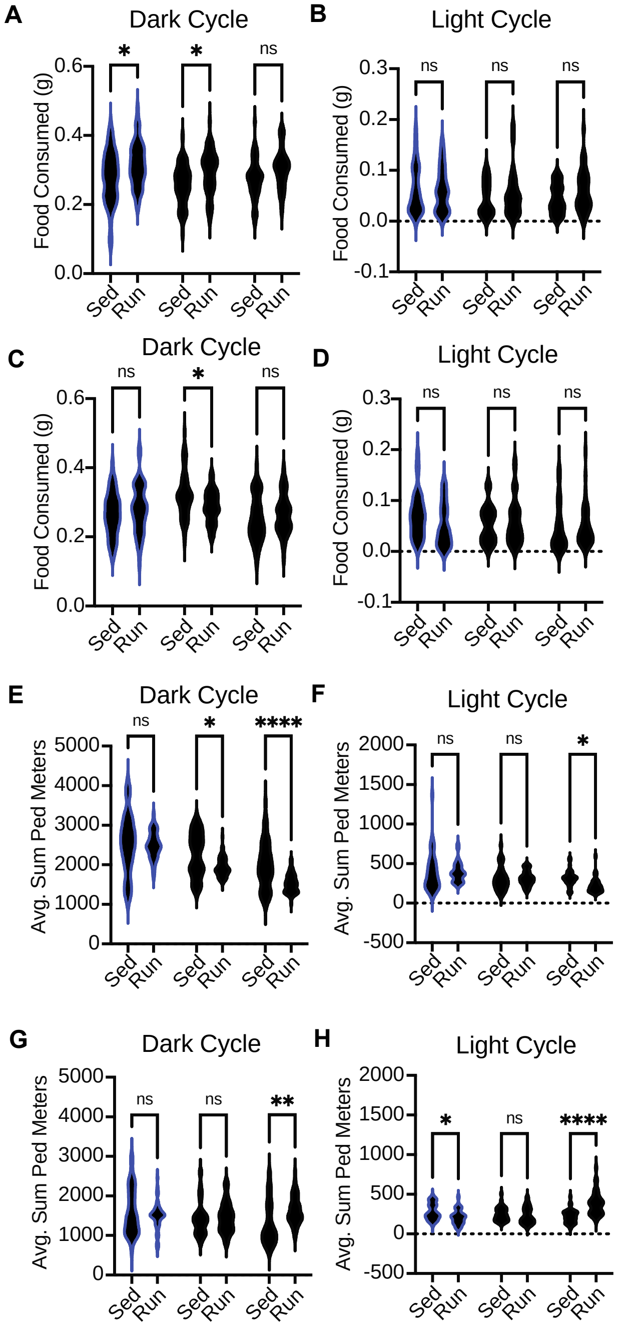


**Supp Fig. 7**

**(A-B)** Bonferroni posthoc results for food consumption for females during the dark cycle (A) and light cycle (B). **(C-D)** Bonferroni posthoc results for food consumption for males during the dark cycle (C) and light cycle (D). **(E-F)** Bonferroni posthoc results for the average sum of ped meters n for females during the dark cycle (E) and light cycle (F). **(G-H)** Bonferroni posthoc results for the average sum of ped meters for males during the dark cycle (G) and light cycle (H). Thick white line indicates median, thin white lines indicate quartiles. *P < 0.05, **P < 0.01, ***P < 0.001, ****P < 0.0001.

**Supp Fig. 8**

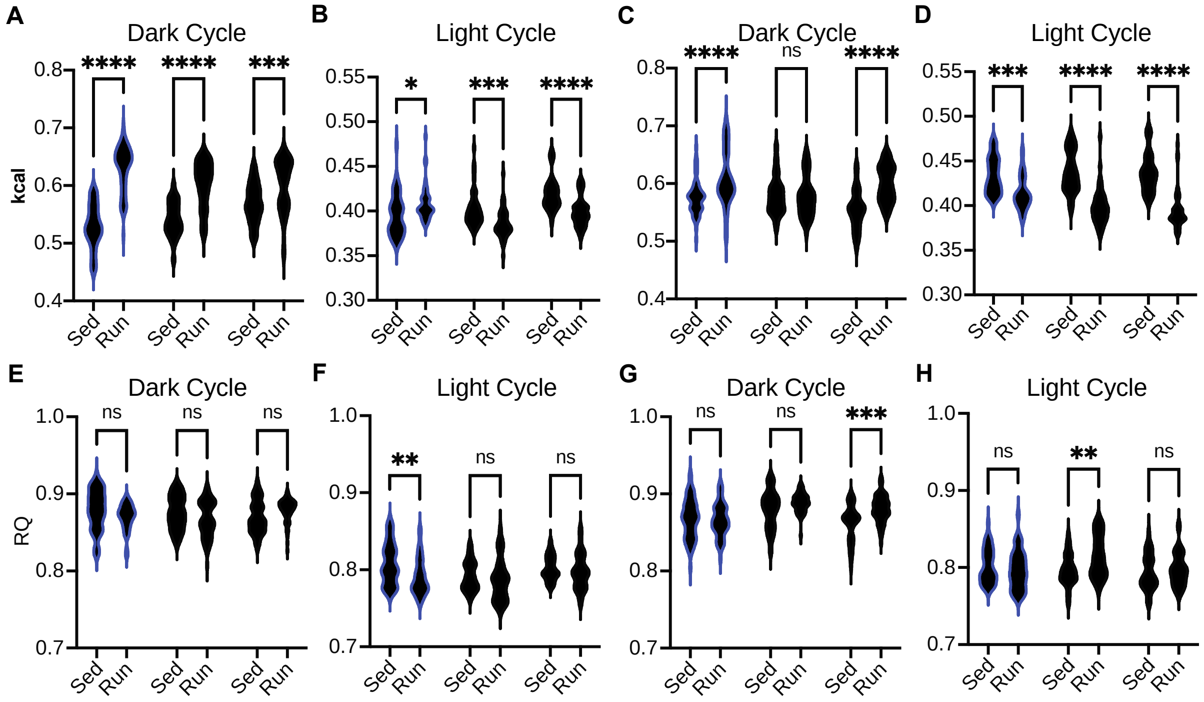


**Supp Fig. 8**

**(A-B)** Bonferroni posthoc results for energy expenditure (kcal/hr) for females during the dark cycle (A) and light cycle (B). **(C-D)** Bonferroni posthoc results for energy expenditure (kcal/hr) for males during the dark cycle (C) and light cycle (D). **(E-F)** Bonferroni posthoc results for RQ for females during the dark cycle (E) and light cycle (F). **(G-H)** Bonferroni posthoc results for RQ for males during the dark cycle (G) and light cycle (H).

**Supp Fig. 9**

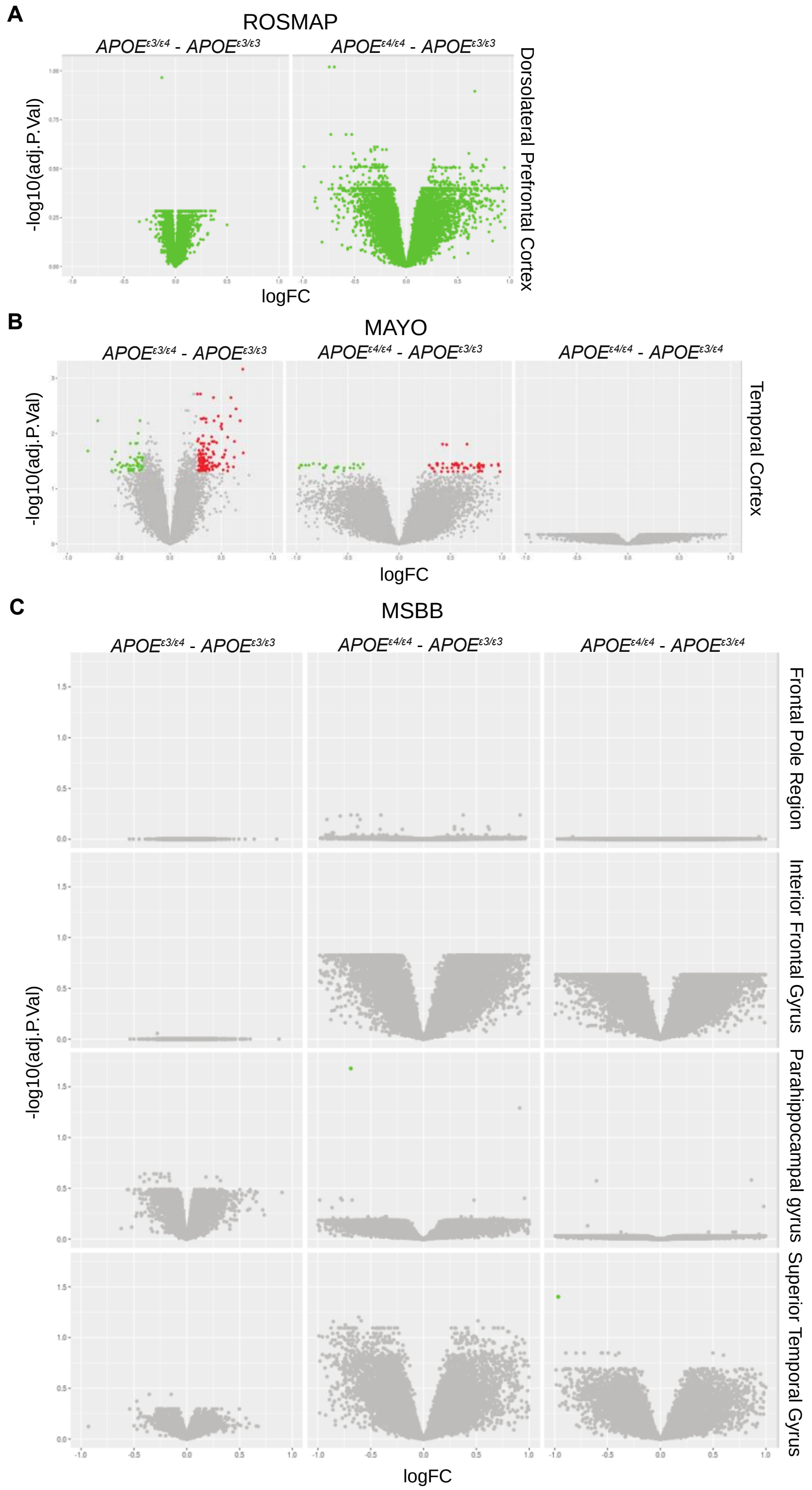


**Supp Fig. 9**
**(A)** ROSMAP (syn8456629) volcano plots for DE genes (FDR<0.05, logFC>1.2) across APOE genotype comparisons, green indicates not significant. **(B)** MAYO (syn8466812) volcano plots

for DE genes (FDR<=0.05) across APOE genotype comparisons, green indicates significantly downregulated, grey indicates not significant, red indicates significantly upregulated.
**(C)** Mount Sinai Brain Bank (MSBB) (syn8484987) volcano plots for DE genes (FDR<=0.05) across APOE genotype comparisons, green indicates significantly downregulated, grey indicates not significant.

**Supp Fig. 10**

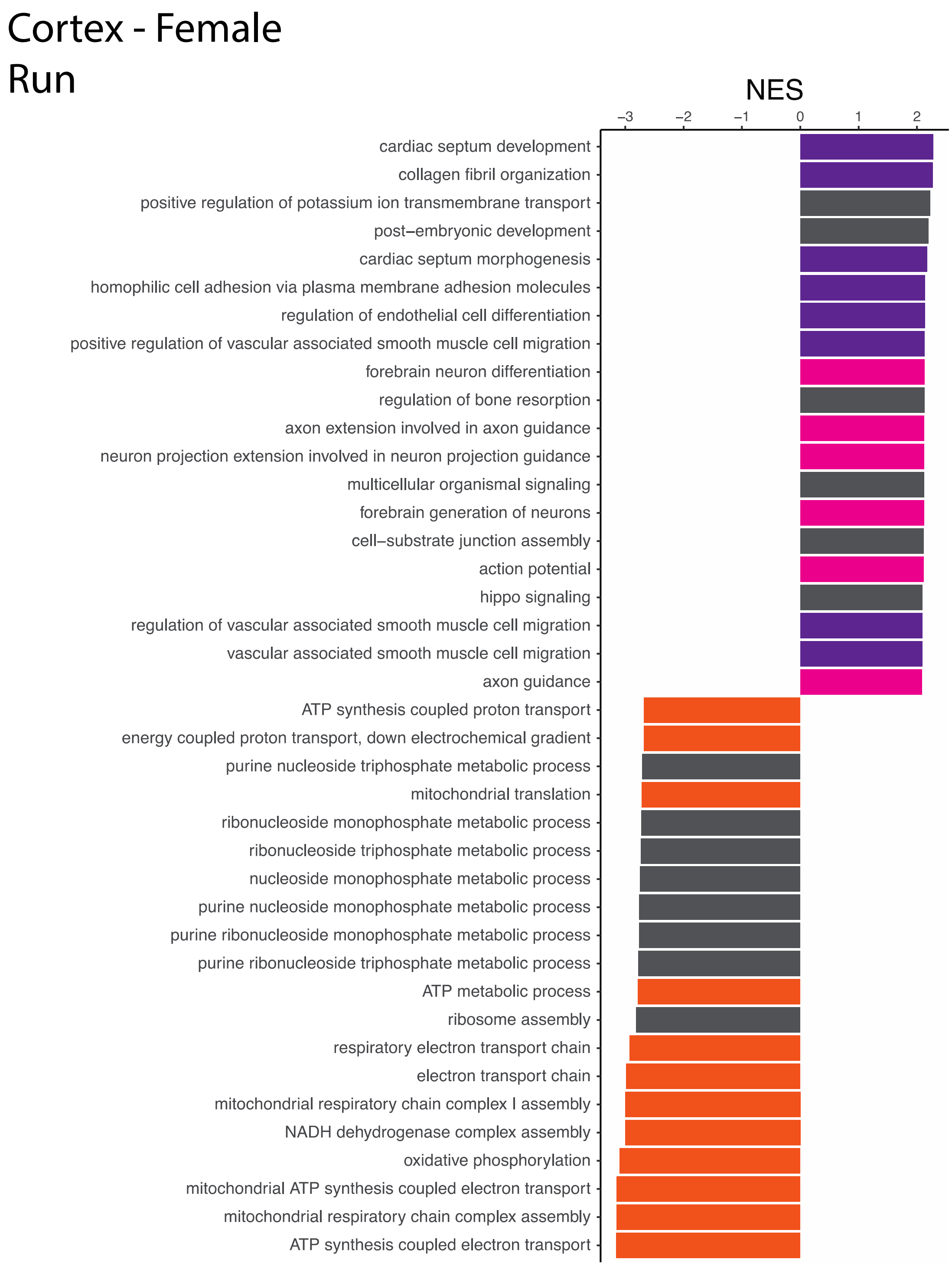


**Supp Fig. 10**
Highest and lowest Normalized Enrichment Scores (NES) for Female Cortex for the main effect of Running. Highlights include: purple = vascular integrity, pink = neuronal/synaptic health, yellow = cellular motility, orange = mitochondrial metabolism, green = immune system response, grey = other.

**Supp Fig. 11**

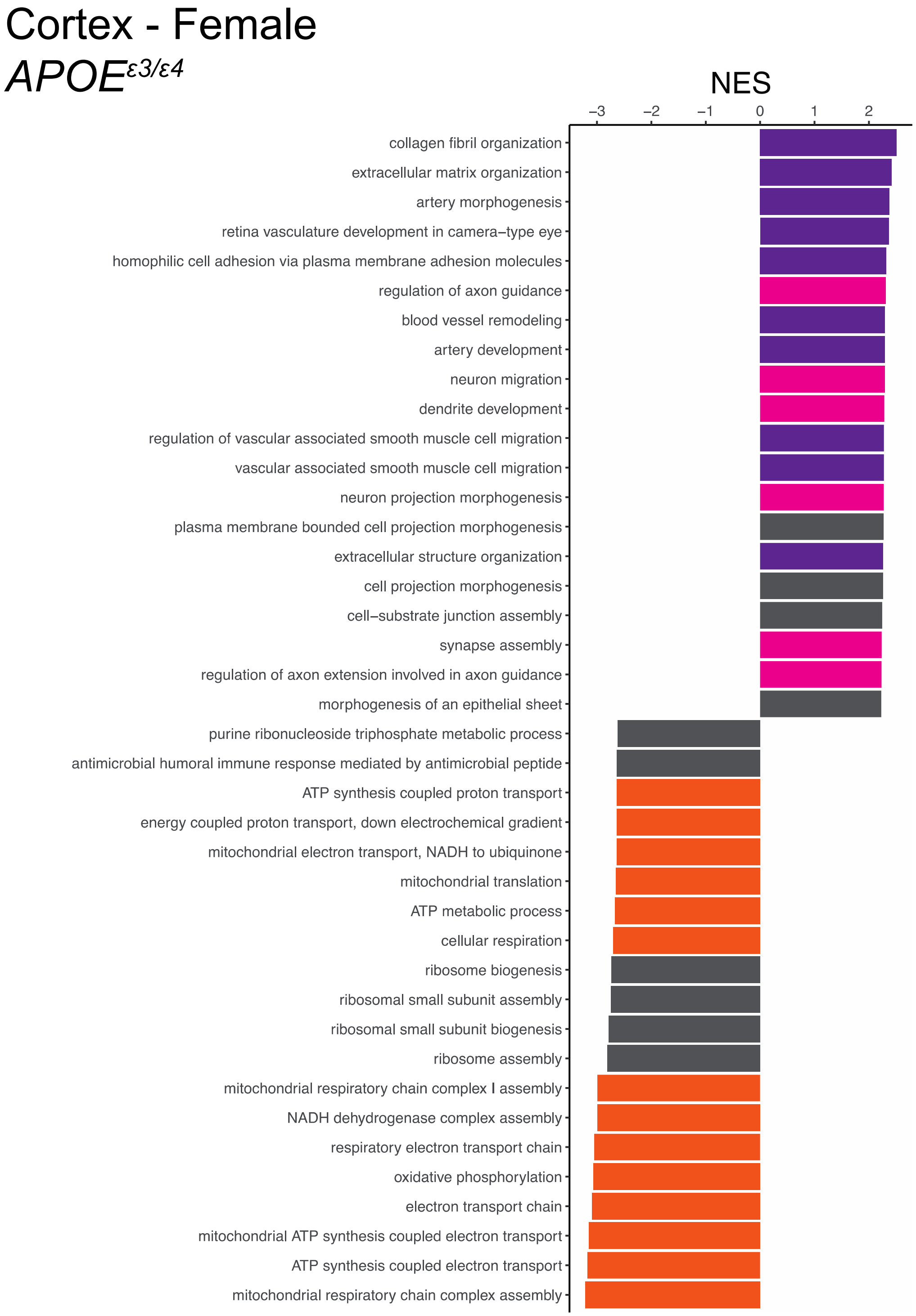


**Supp Fig. 11**
Highest and lowest Normalized Enrichment Scores (NES) for Female Cortex for the main effect of *APOE*$^{\varepsilon3/\varepsilon4}$. Highlights include: purple = vascular integrity, pink = neuronal/synaptic health, yellow = cellular motility, orange = mitochondrial metabolism, green = immune system response, grey = other.

**Supp Fig. 12**

### Cortex - Female

### *APOE*$^{\varepsilon3/\varepsilon4}$:Run

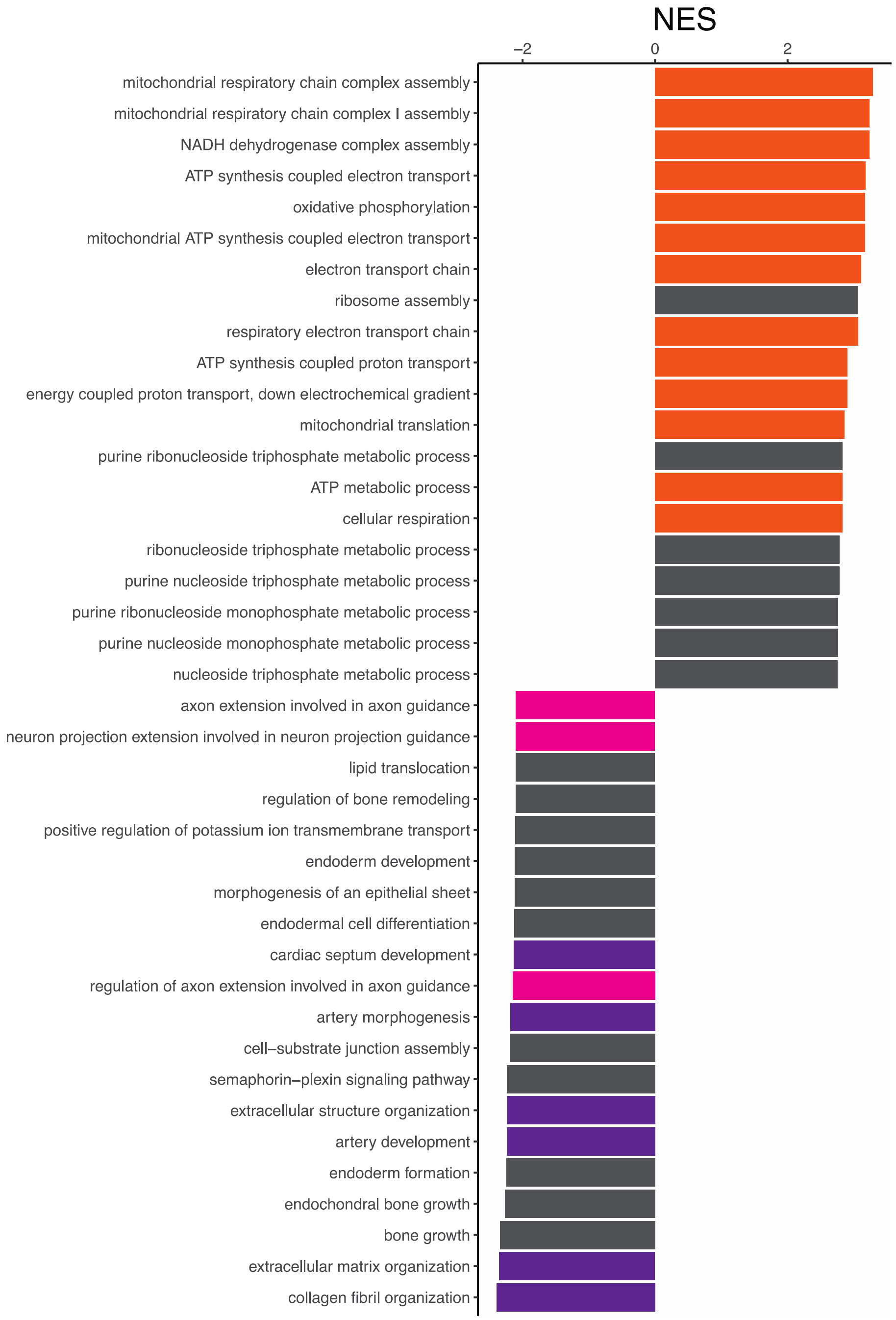


**Supp Fig. 12**
Highest and lowest Normalized Enrichment Scores (NES) for Female Cortex for the interactive effect of *APOE*$^{\varepsilon3/\varepsilon4}$:Run. Highlights include: purple = vascular integrity, pink = neuronal/synaptic health, yellow = cellular motility, orange = mitochondrial metabolism, green = immune system response, grey = other.

**Supp Fig. 13**

### Cortex - Female
#### *APOE*$^{\varepsilon4/\varepsilon4}$

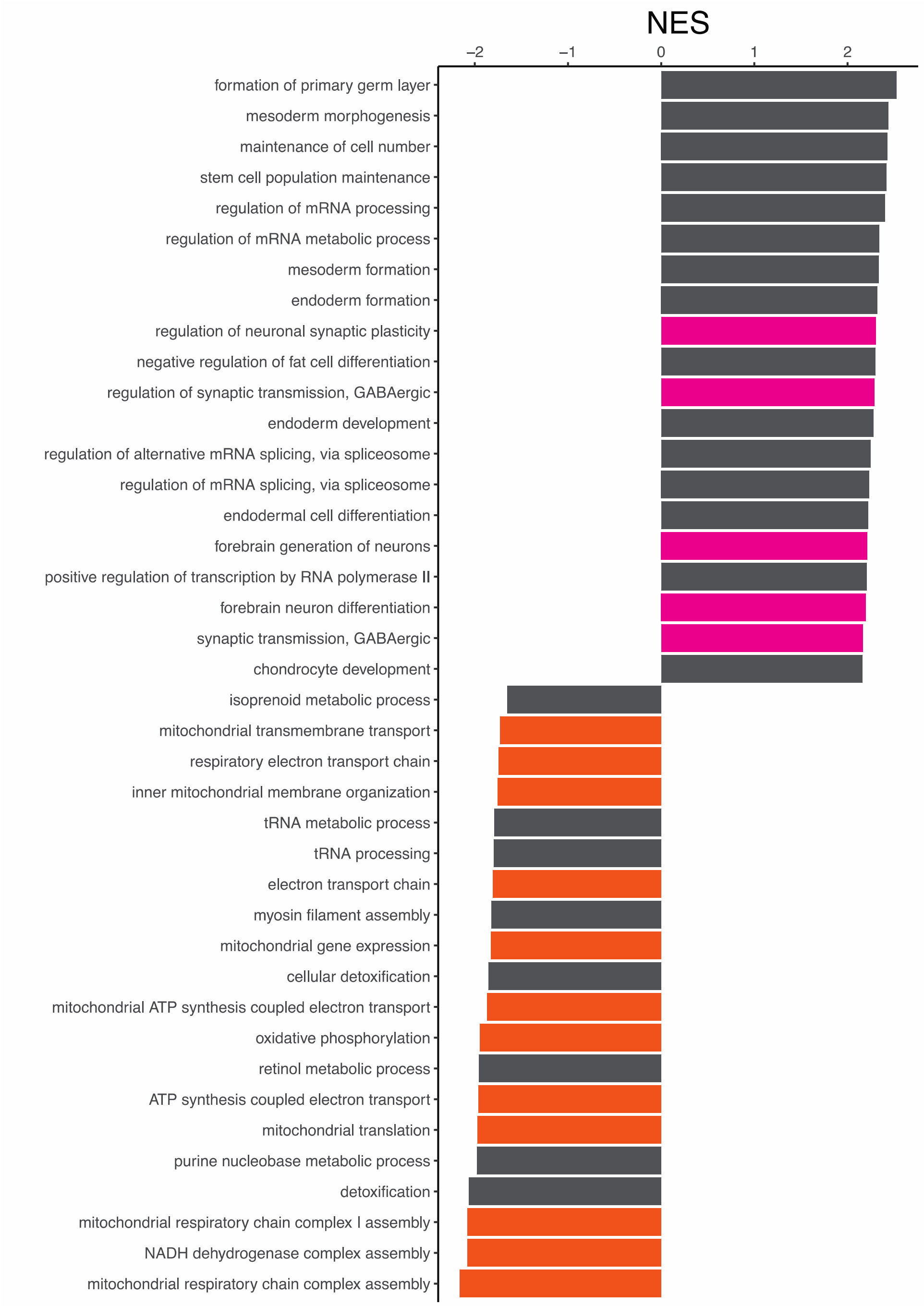


**Supp Fig. 13**
Highest and lowest Normalized Enrichment Scores (NES) for Female Cortex for the main effect of *APOE*$^{\varepsilon4/\varepsilon4}$. Highlights include: purple = vascular integrity, pink = neuronal/synaptic health, yellow = cellular motility, orange = mitochondrial metabolism, green = immune system response, grey = other.

**Supp Fig. 14**

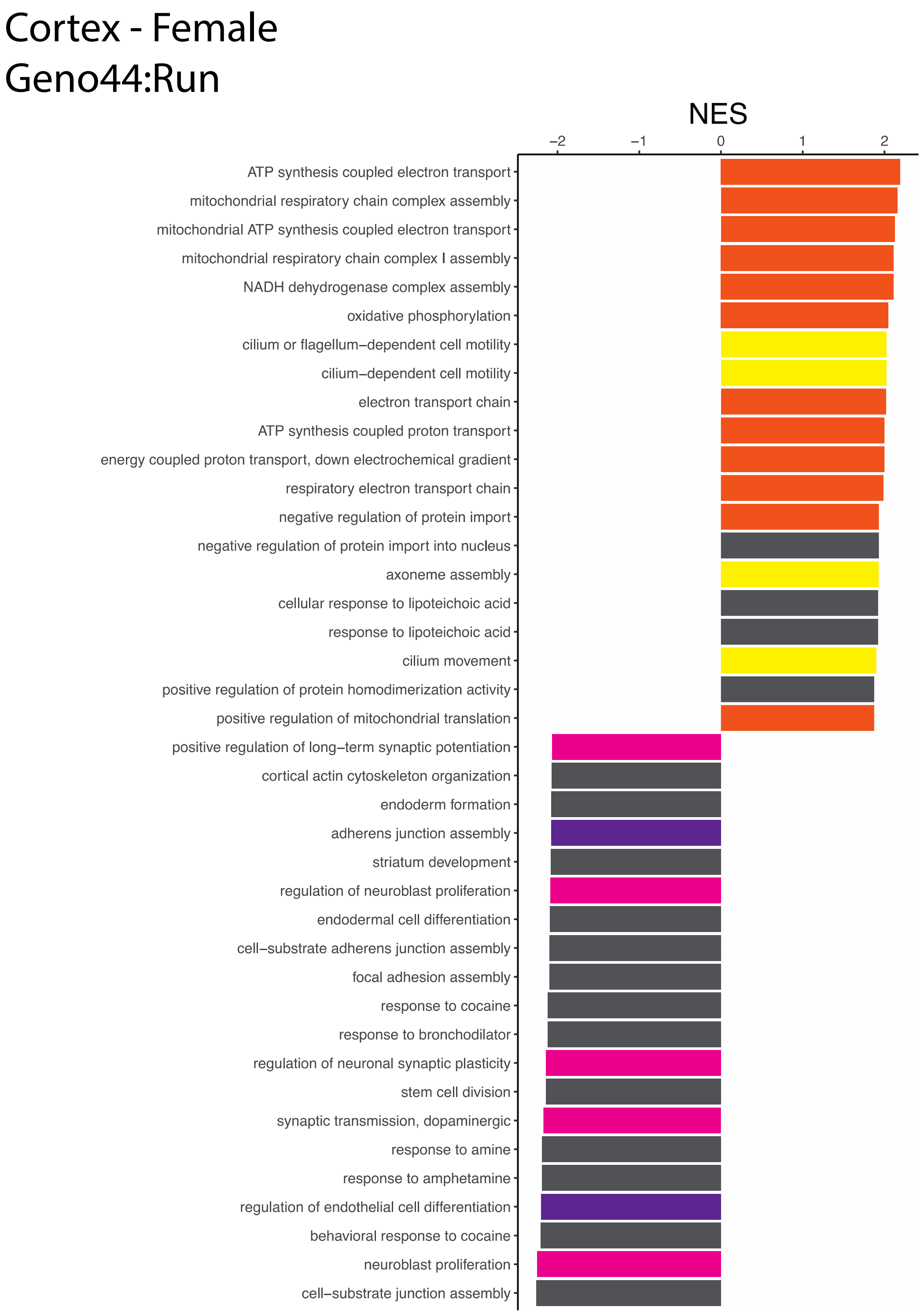


**Supp Fig. 14**
Highest and lowest Normalized Enrichment Scores (NES) for Female Cortex for the interactive effect of *APOE*$^{\varepsilon 4/\varepsilon 4}$:Run. Highlights include: purple = vascular integrity, pink = neuronal/synaptic health, yellow = cellular motility, orange = mitochondrial metabolism, green = immune system response, grey = other.

**Supp Fig. 15**

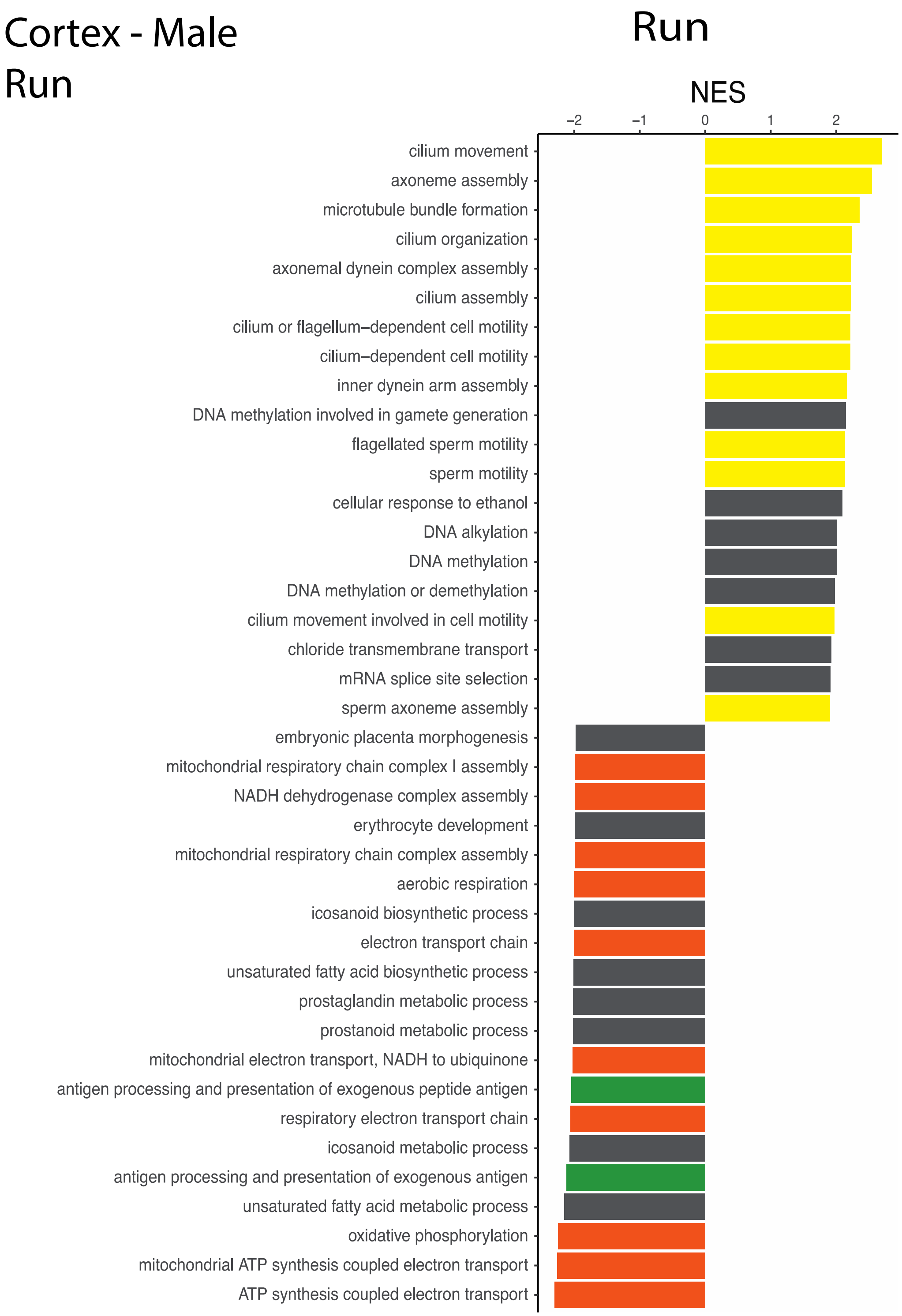


**Supp Fig. 15**
Highest and lowest Normalized Enrichment Scores (NES) for Male Cortex for the main effect of Running. Highlights include: purple = vascular integrity, pink = neuronal/synaptic health, yellow = cellular motility, orange = mitochondrial metabolism, green = immune system response, grey = other.

**Supp Fig. 16**

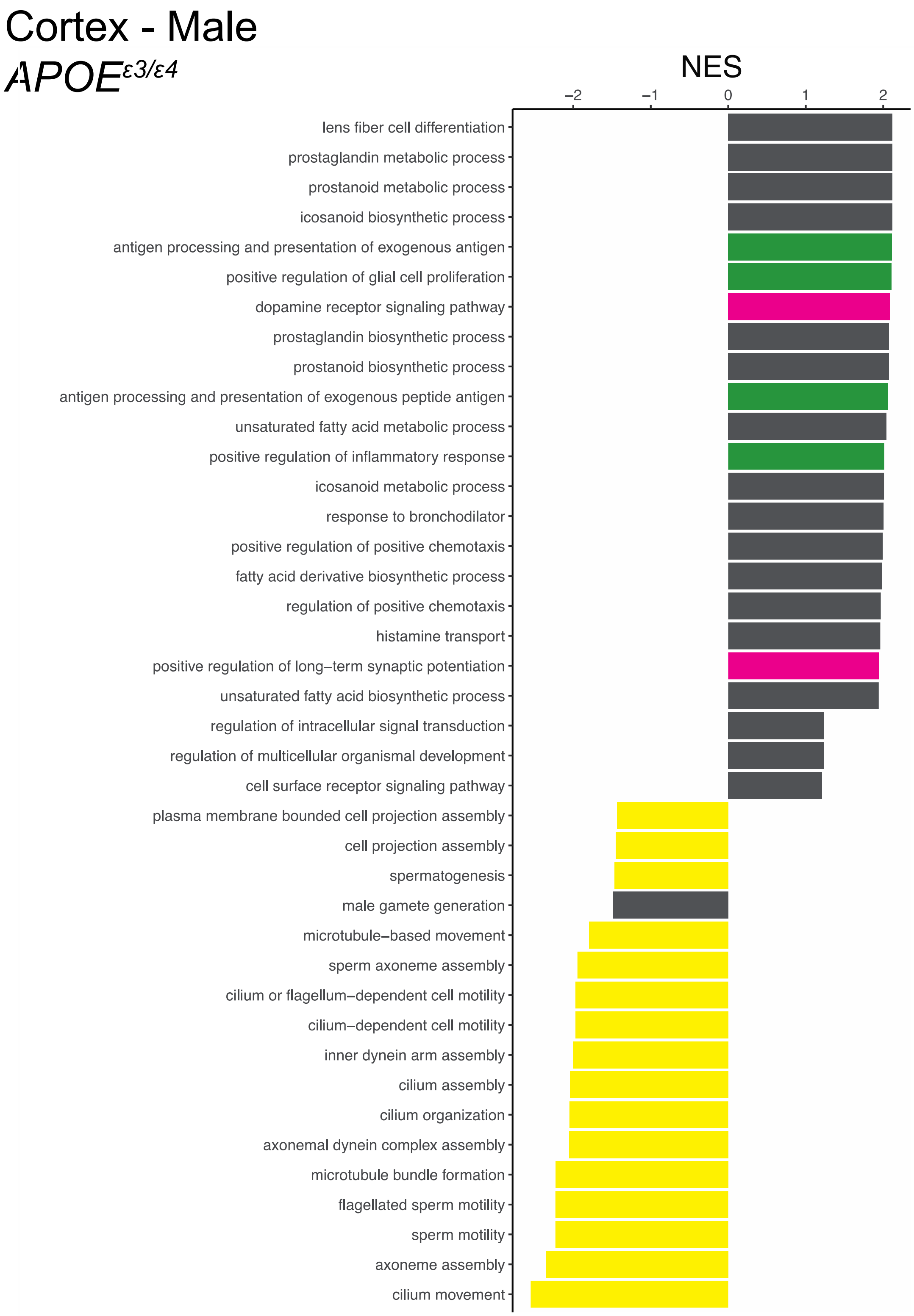


**Supp Fig. 16**
Highest and lowest Normalized Enrichment Scores (NES) for Male Cortex for the main effect of *APOE*$^{\varepsilon3/\varepsilon4}$. Highlights include: purple = vascular integrity, pink = neuronal/synaptic health, yellow = cellular motility, orange = mitochondrial metabolism, green = immune system response, grey = other.

**Supp Fig. 17**

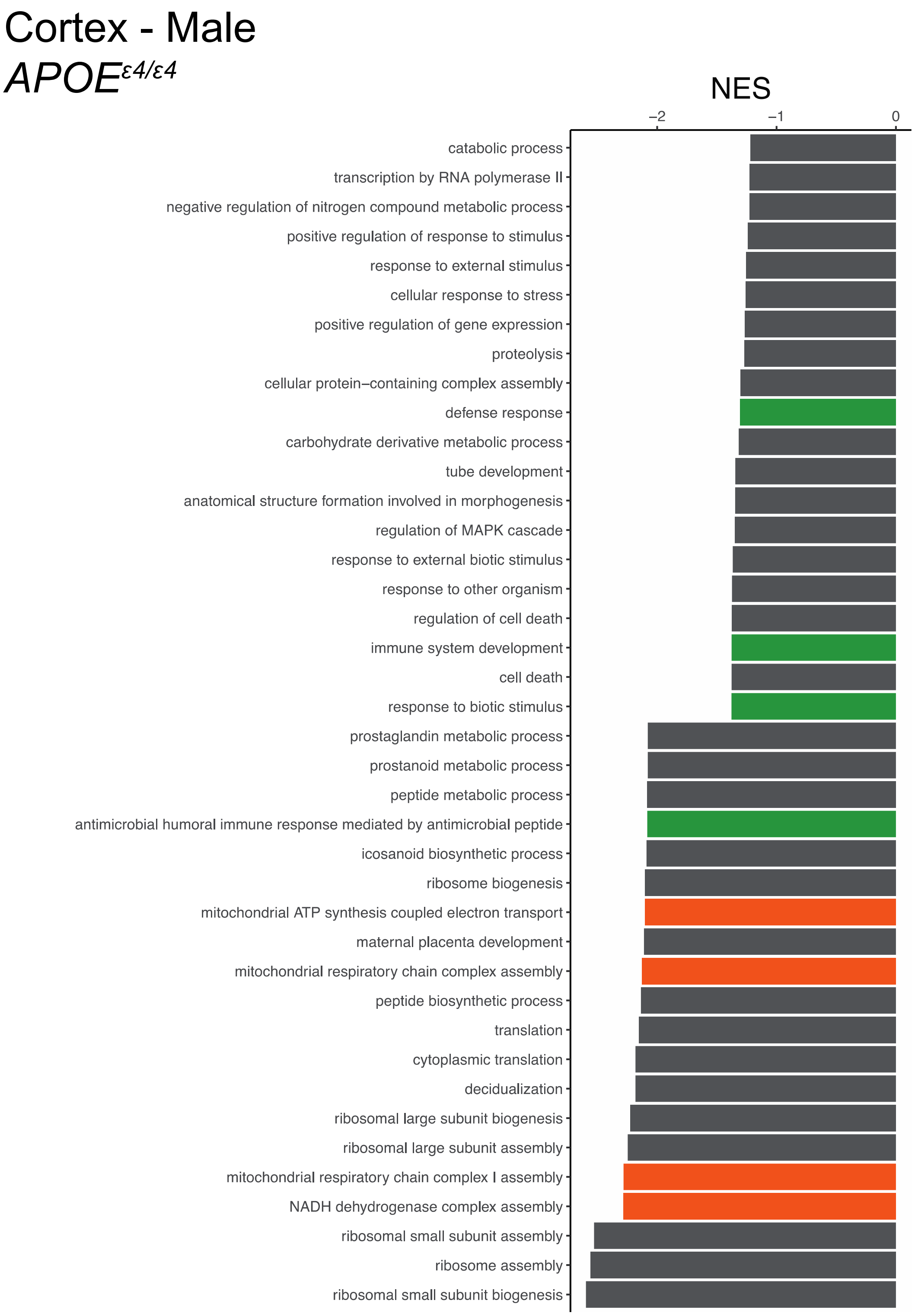


**Supp Fig. 17**
Highest and lowest Normalized Enrichment Scores (NES) for Male Cortex for the main effect of *APOE*$^{\varepsilon4/\varepsilon4}$. Highlights include: purple = vascular integrity, pink = neuronal/synaptic health, yellow = cellular motility, orange = mitochondrial metabolism, green = immune system response, grey = other.

**Supp Fig. 18**

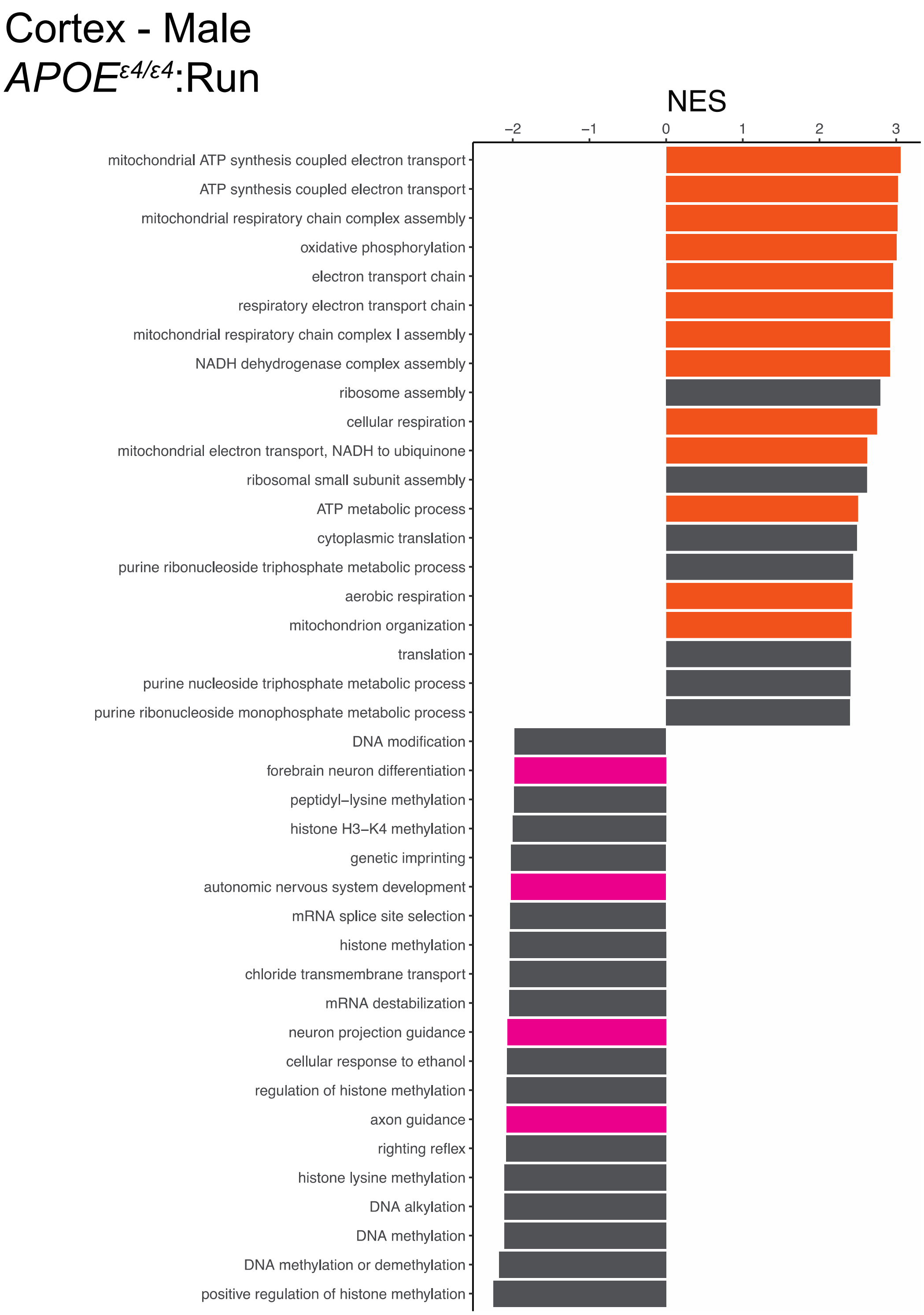


**Supp Fig. 18**
Highest and lowest Normalized Enrichment Scores (NES) for Male Cortex for the interactive effect of *APOE*$^{\varepsilon 4/\varepsilon 4}$:Run. Highlights include: purple = vascular integrity, pink = neuronal/synaptic health, yellow = cellular motility, orange = mitochondrial metabolism, green = immune system response, grey = other.

**Supp Fig. 19**

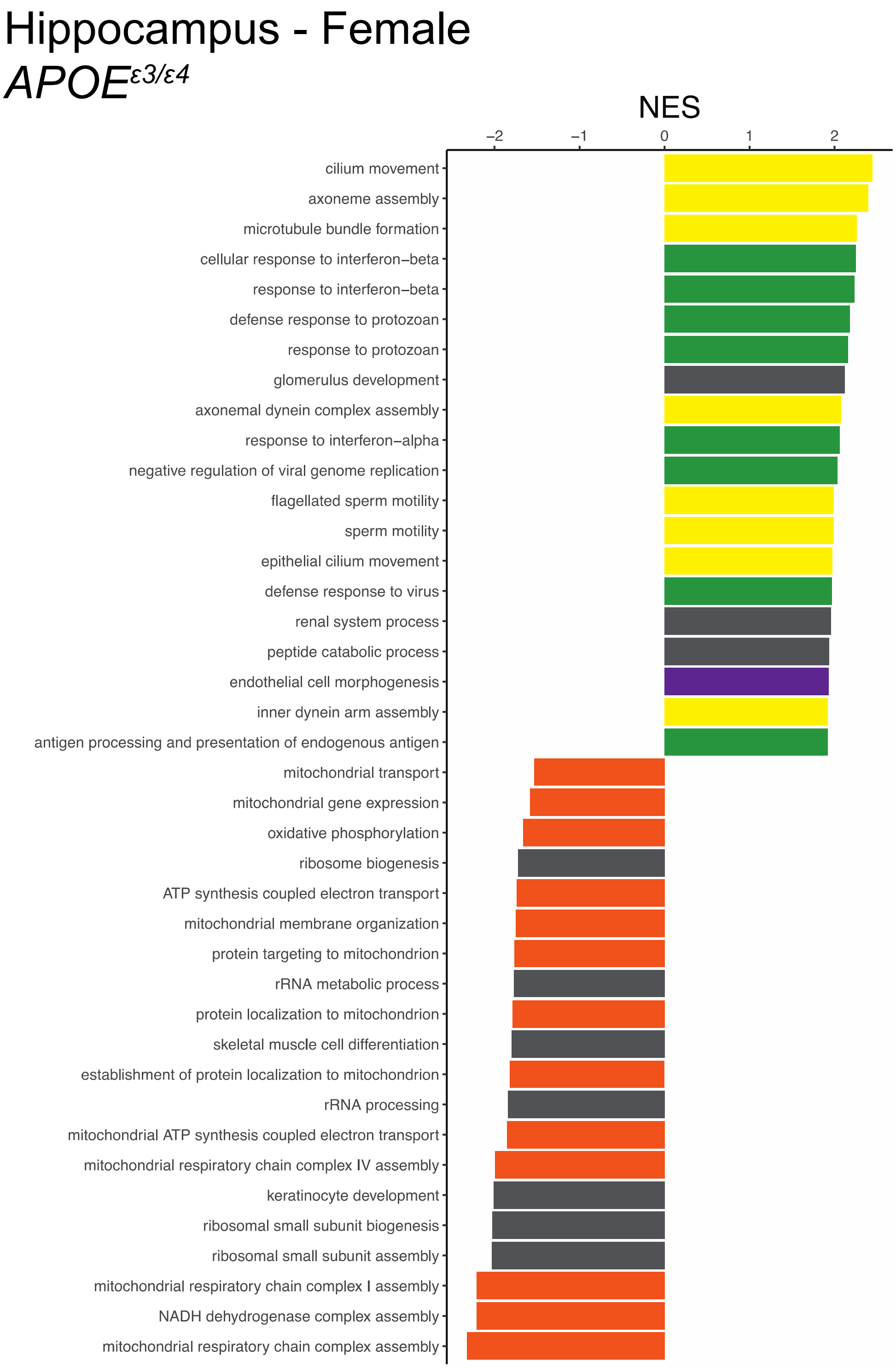


**Supp Fig. 19**
Highest and lowest Normalized Enrichment Scores (NES) for Female Hippocampus for the main effect of *APOE*$^{\varepsilon3/\varepsilon4}$. Highlights include: purple = vascular integrity, pink = neuronal/synaptic health, yellow = cellular motility, orange = mitochondrial metabolism, green = immune system response, grey = other.

**Supp Fig. 20**

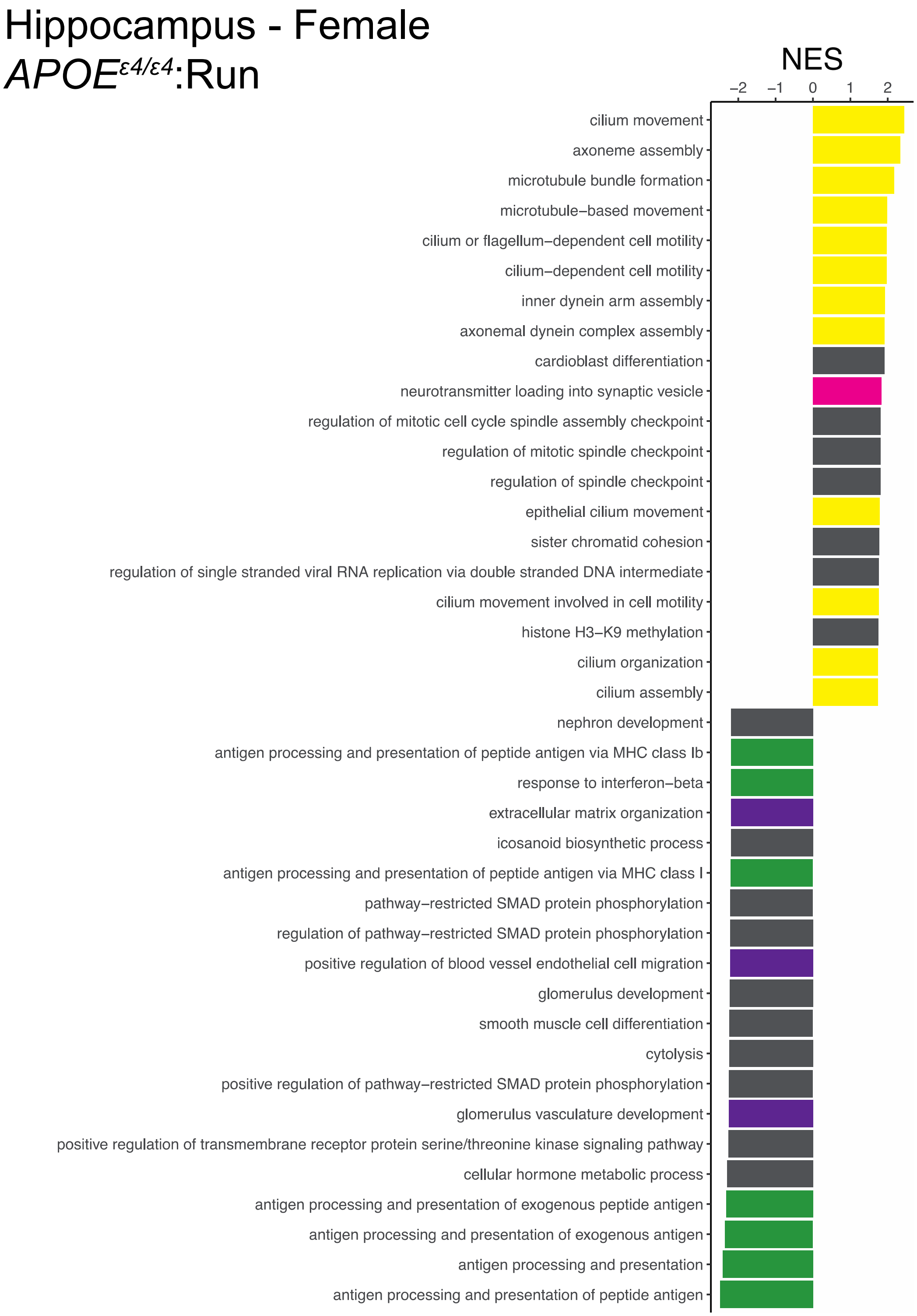


**Supp Fig. 20**
Highest and lowest Normalized Enrichment Scores (NES) for Female Hippocampus for the interactive effect of *APOE*$^{ε3/ε4}$:Run. Highlights include: purple = vascular integrity, pink = neuronal/synaptic health, yellow = cellular motility, orange = mitochondrial metabolism, green = immune system response, grey = other.

**Supp Fig. 21**

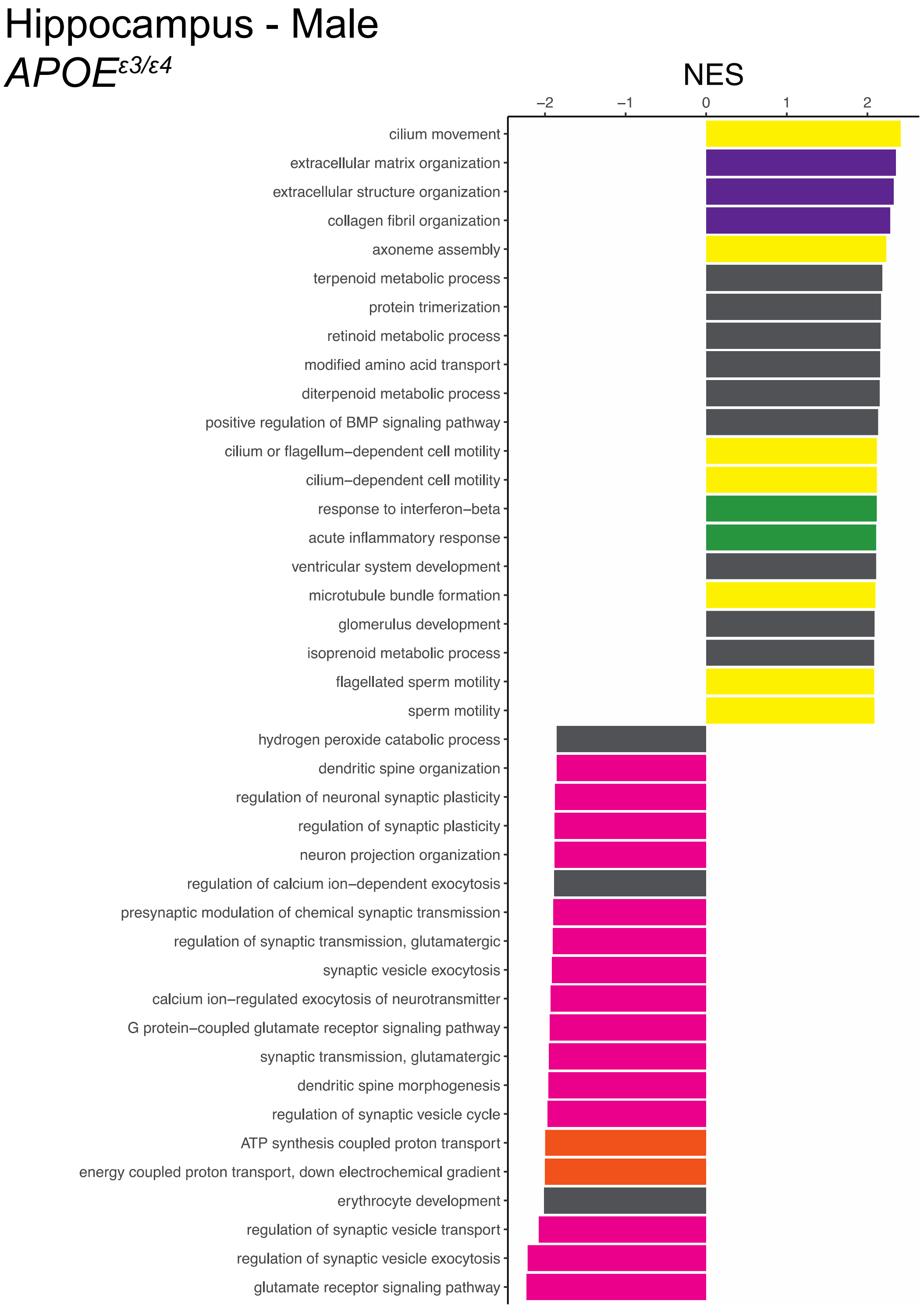


**Supp Fig. 21**
Highest and lowest Normalized Enrichment Scores (NES) for Male Hippocampus for the main effect of *APOE*$^{\varepsilon 3/\varepsilon 4}$. Highlights include: purple = vascular integrity, pink = neuronal/synaptic health, yellow = cellular motility, orange = mitochondrial metabolism, green = immune system response, grey = other.

**Supp Fig. 22**

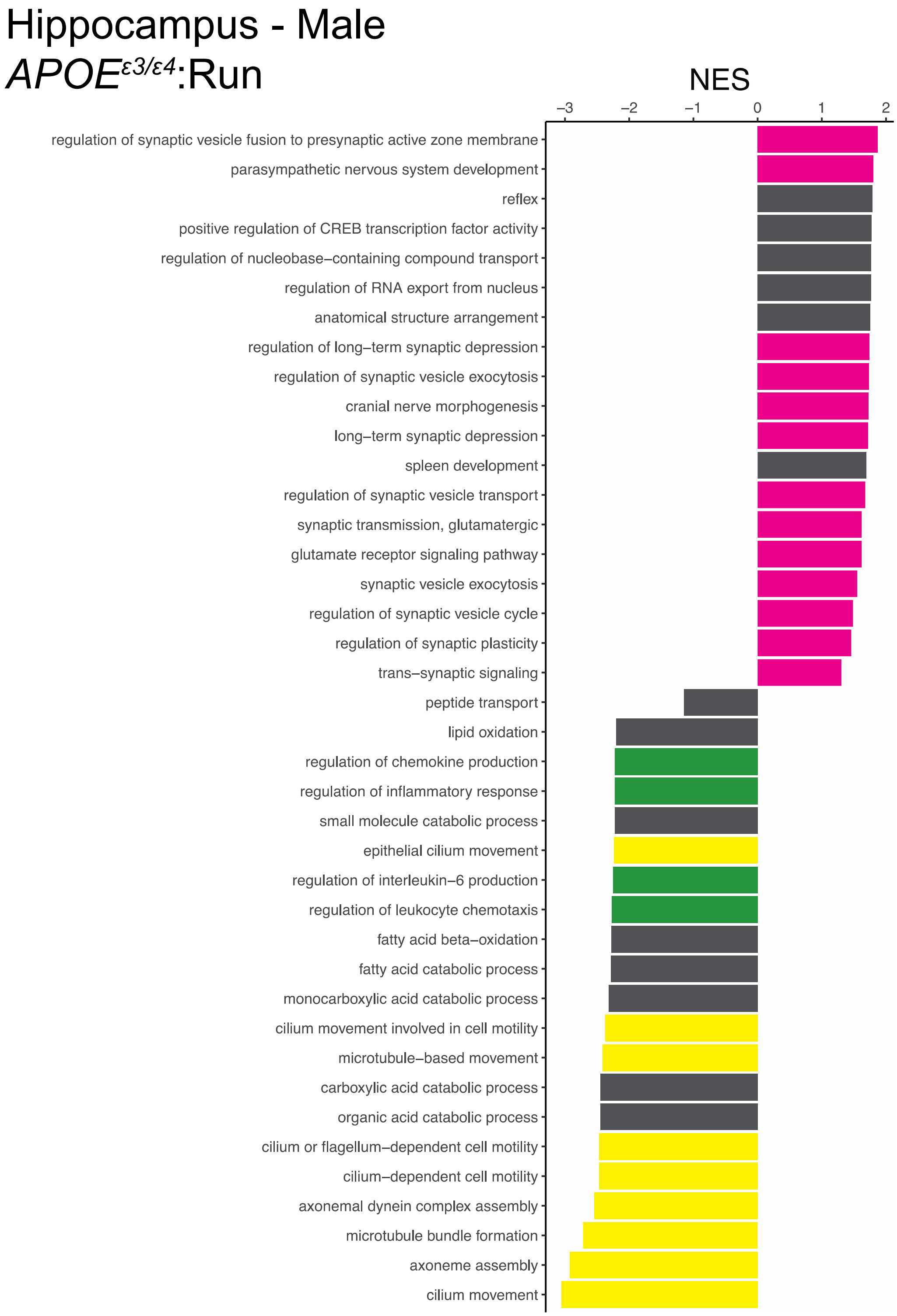


**Supp Fig. 22**
Highest and lowest Normalized Enrichment Scores (NES) for Male Hippocampus for the interactive effect of *APOE*$^{\varepsilon 3/\varepsilon 4}$:Run. Highlights include: purple = vascular integrity, pink = neuronal/synaptic health, yellow = cellular motility, orange = mitochondrial metabolism, green = immune system response, grey = other.

**Supp Fig. 23**

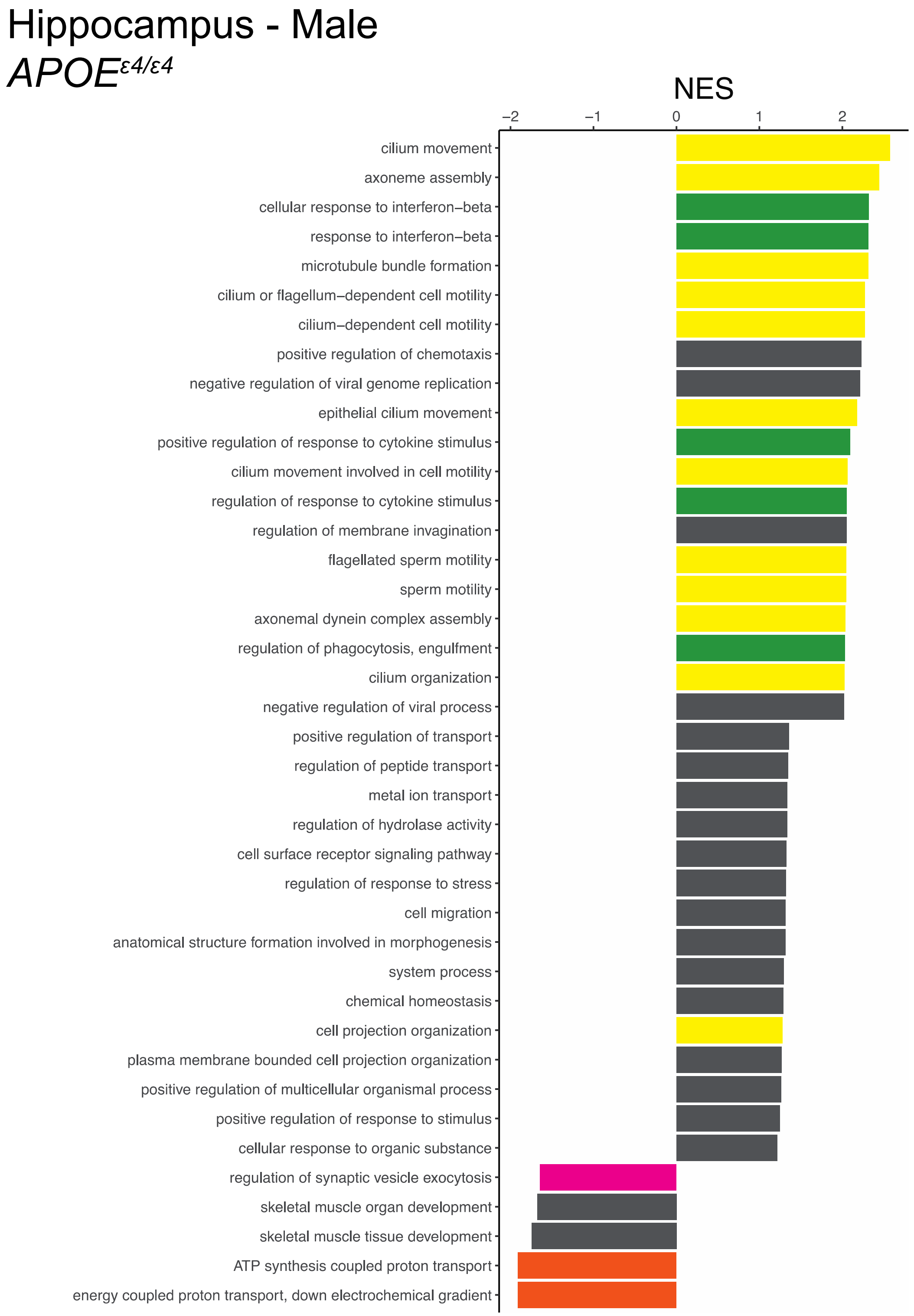


**Supp Fig. 23**

Highest and lowest Normalized Enrichment Scores (NES) for Male Hippocampus for the interactive effect of *APOE*$^{\varepsilon 4/\varepsilon 4}$. Highlights include: purple = vascular integrity, pink = neuronal/synaptic health, yellow = cellular motility, orange = mitochondrial metabolism, green = immune system response, grey = other.

**Supp Fig. 24**

### Hippocampus - Male

### *APOE*$^{\varepsilon4/\varepsilon4}$:Run

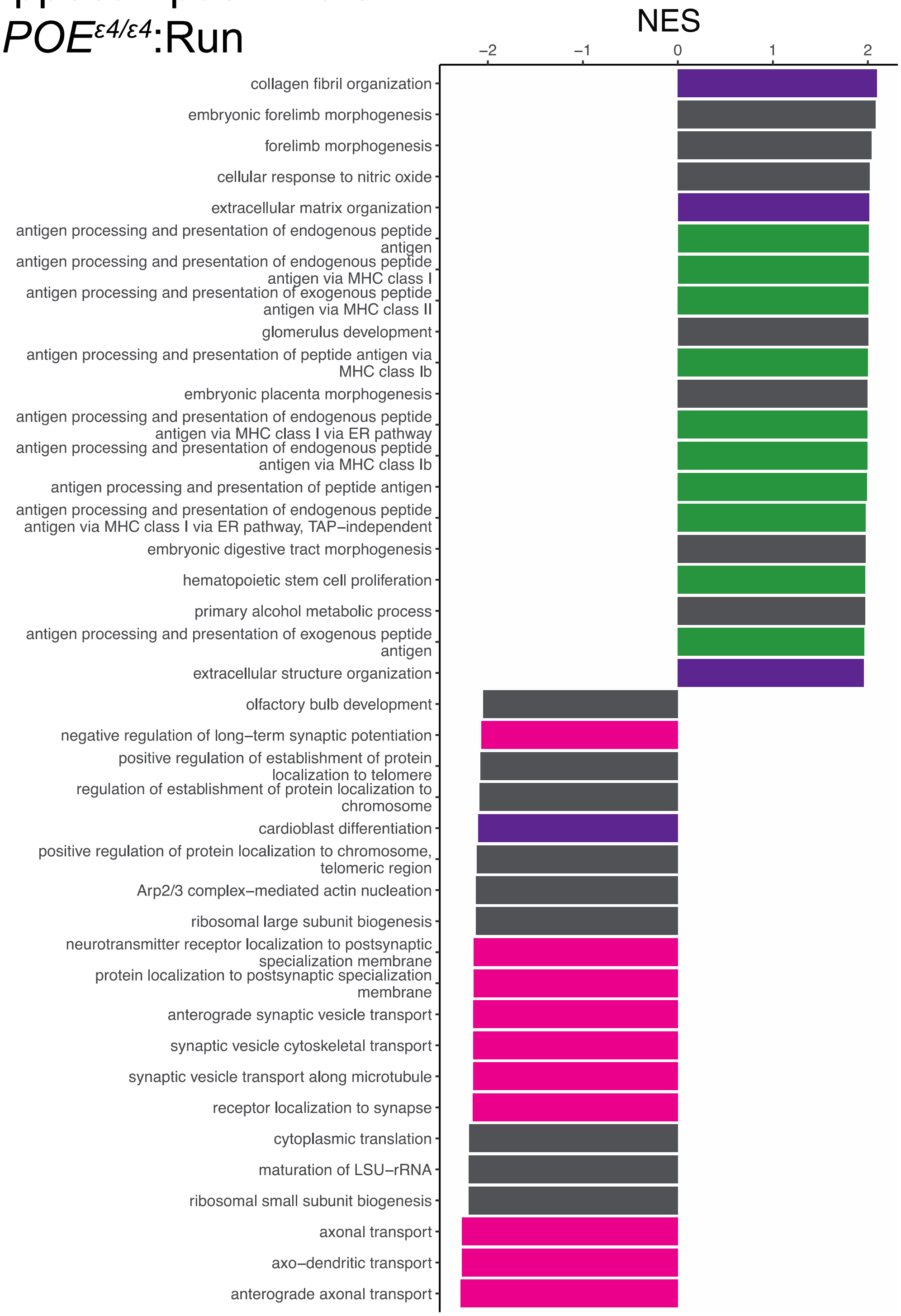


**Supp Fig. 24**

Highest and lowest Normalized Enrichment Scores (NES) for Male Hippocampus for the interactive effect of *APOE*$^{\varepsilon4/\varepsilon}$:Run. Highlights include: purple = vascular integrity, pink = neuronal/synaptic health, yellow = cellular motility, orange = mitochondrial metabolism, green = immune system response, grey = other.
